# DeepBryo: a web app for AI-assisted morphometric characterization of cheilostome bryozoans

**DOI:** 10.1101/2022.11.17.516938

**Authors:** Emanuela Di Martino, Björn Berning, Dennis P Gordon, Piotr Kuklinski, Lee Hsiang Liow, Mali H Ramsfjell, Henrique L Ribeiro, Abigail M Smith, Paul D Taylor, Kjetil L Voje, Andrea Waeschenbach, Arthur Porto

## Abstract

1. Bryozoans are becoming an increasingly popular study system in macroevolutionary, ecological, and paleobiological research. Members of this colonial invertebrate phylum are notable for displaying an exceptional degree of division of labor in the form of specialized modules (polymorphs), which allow for the inference of individual allocation of resources to reproduction, defense, and growth using simple morphometric tools. However, morphometric characterizations of bryozoans are notoriously labored, due to the high number of structures often captured per image, as well as the need for specialized knowledge necessary for classifying individual skeletal structures within those images.
2. We here introduce DeepBryo, a web application for deep learning-based morphometric characterization of cheilostome bryozoans. DeepBryo requires a single image as input and performs measurements automatically using instance segmentation algorithms. DeepBryo is capable of detecting objects belonging to six classes and outputting fourteen morphological shape measurements for each object based on the inferred segmentation maps. The users can visualize the predictions, check for errors, and directly filter model outputs on the web browser. Measurements can then be downloaded as a comma-separated values file.
3. DeepBryo has been trained and validated on a total of 72,412 structures, belonging to six different object classes in 935 SEM images of cheilostome bryozoans belonging to 109 different families. The model shows high (>0.8) recall and precision for zooid-level structures. Its misclassification rate is low (~4%) and largely concentrated in a single object class (opesia). The model’s estimated structure-level area, height, and width measurements are statistically indistinguishable from those obtained via manual annotation (r^2^ varying from 0.89 to 0.98) and show no detectable bias. DeepBryo reduces the person-hours required for characterizing the zooids in individual colonies to less than 1% of the time required for manual annotation at no significant loss of measurement accuracy.
4. Our results indicate that DeepBryo enables cost-, labor,- and time-efficient morphometric characterization of cheilostome bryozoans. DeepBryo can greatly increase the scale of macroevolutionary, ecological, taxonomic, and paleobiological analyses, as well as the accessibility of deep learning tools for this emerging model system. Finally, DeepBryo provides the building blocks necessary for adapting the current application to other study groups.

## 1 Introduction

Recent advances in macroevolutionary research emphasize that a large number of evolutionary phenomena are better explained when using data from both extant and extinct taxa (Love et al., 2022; Mongiardino Koch et al., 2021; Parry et al., 2016; Slater, 2013; Slater et al., 2012). Yet, most macroevolutionary analyses of trait diversification patterns have been disproportionately focused on organisms with poor fossil records (e.g., terrestrial vertebrates and plants) (Orr et al., 2022). As a consequence, our understanding of trait diversification dynamics is still highly biased by the limited temporal sampling of most datasets currently being used to infer trait dynamics, as well as the difficulties of integrating neontological and paleontological datasets into a single phylogenetic framework. Compounding this bias, most empirical studies of trait diversification focus on one trait at a time and are therefore blind to the multivariate nature of genotype-to-phenotype maps, in which pleiotropy, epistasis, and genotype-by-environment interactions are prevalent (Houle et al., 2010; Melo et al., 2016). This univariate focus of macroevolutionary research is not surprising when taking into account the labor and financial costs associated with high-dimensional phenotyping. It does imply that our current views of macroevolutionary diversification often do not account for the role of trait correlations (Melo et al., 2016).

Members of the colonial phylum Bryozoa have played a fundamental role in classical debates regarding tempo and mode of morphological evolution (Cheetham, 1986a, 1986b, 1987; Cheetham et al., 1993, 1994; Voje et al., 2020), and represent an emerging model system in macroevolutionary, ecological, and paleobiological research (Clark et al., 2010; Di Martino & Liow, 2021, 2022; Liow et al., 2017; Liow & Taylor, 2019; O’Dea & Jackson, 2009, 2002; O’Dea & Okamura, 2000; Okamura et al., 2013). Bryozoans have an old (Cambrian) (Zhang et al., 2021), species-rich (>22,000 described species), and high temporal resolution fossil record (Taylor, 2020). Their skeletal features are finely preserved over large timescales, to the point where fossil taxa can often be identified at the species level (Jackson & Cheetham, 1990; Simpson & Jackson, 2022). Order Cheilostomatida, in particular, represents the most abundant and diverse order within Bryozoa, containing around 80% of the phylum’s living species diversity plus as many as 7,900 described fossil species (Bock & Gordon, 2013). Members of this order are known for the high degree of modularity (and division of labor) in their colonies, in which genetically identical zooids can develop into a wide array of polymorphs (Schack et al., 2019). These zooid polymorphs and modified skeletal structures can devote themselves to functions such as reproduction (ovicells), defense (avicularia), feeding (autozooid), structural integrity (kenozooids), among others, and provide a window into how individual colonies allocate resources in response to environmental cues (Schack et al., 2019), given their genetic background.

For a long time (e.g, Buss, 1979), researchers have emphasized that the biology of bryozoans provides a unique opportunity to study ecological and life-history traits in the fossil record, in which colony-level allocation of resources can be estimated using simple morphometric tools (Di Martino & Liow, 2021; Liow et al., 2017; O’Dea, 2003; O’Dea & Jackson, 2002; O’Dea & Okamura, 2000). On the analytical front, the main bottleneck for a more widespread adoption of bryozoans as a model system in macroevolutionary biology is the laborious nature of the morphometric data. In particular, bryozoan skeletal morphology is usually captured using scanning electron microscopy (SEM) images of dry specimens, which can contain hundreds of structures within just one image. The recognition and classification of these structures are not trivial and often require expert domain knowledge (see (Schack et al., 2019)). Morphometric characterizations of skeletal morphology in the group are thus usually carried out manually by a dwindling number of taxonomic experts, a process that is low-throughput, costly, and time-consuming to reproduce.

Over the last decade, rapid changes in morphometric investigations have been spurred by developments in computer vision and machine learning (Lürig et al., 2021). In particular, deep-learning-based computer vision tools have shown exceptional performance in a wide variety of tasks both in scientific and real-world settings (e.g., (Lin et al., 2014)). Consequently, vision-based morphometric approaches are fast becoming an integral part of a biologist’s toolkit, both in the field and the laboratory (Lürig et al., 2021; Porto et al., 2021; Porto & Voje, 2020; Vandaele et al., 2018).

We here introduce DeepBryo, a deep-learning-based web application that allows rapid and accurate morphometric characterization of cheilostome bryozoan images directly in a web browser. DeepBryo approaches the problem of instance segmentation of SEM images using state-of-the-art transformer models (Liu et al., 2021) wrapped in a simple user interface. After uploading an image, the application, the user can visualize, filter, and download segmentation results. Given the recent explosion in the availability of cheilostome SEM images, we expect this simple framework will greatly increase the scale and robustness of macroevolutionary, ecological, taxonomic, and paleobiological research based on this emerging study system.

## 2 DEEPBRYO APP

### 2.1 Description

DeepBryo (Figure 1) is available as a web application (https://www.deepbryo.ngrok.io) or can be deployed locally (https://www.github.com/agporto/DeepBryo). The web application runs on a central server capable of running multiple parallel user sessions using a 12GB NVIDIA QUADRO P4000 GPU. A single SEM image is expected as input. The specimen should lie flat within the image, with its surface perpendicular to the observer. The optimal range of zoom magnification for the input images lies between 30x and 90x, as typically used for bryozoan species identification. Higher or lower magnification will likely result in poorer model predictions and a significant reduction in model recall and precision. Once uploaded, the model predictions are automatically generated and cached using the *streamlit v.1.10.0* python library. The caching mechanism is a key component of DeepBryo, as it allows multiple users to perform filtering and scaling of model predictions without the need to rerun the initial prediction steps. From the user’s point of view, model predictions appear on the browser as a colored mask, in which the color represents six possible object classes: ascopore, autozooid, avicularium, opesia, orifice, ovicell (Figure 2; See Supplemental File 1 for a glossary of terms). Users also have the option of visualizing the corresponding bounding boxes and identification numbers for each structure on the image using the sidebar (Figure S1).

**Figure 1.**
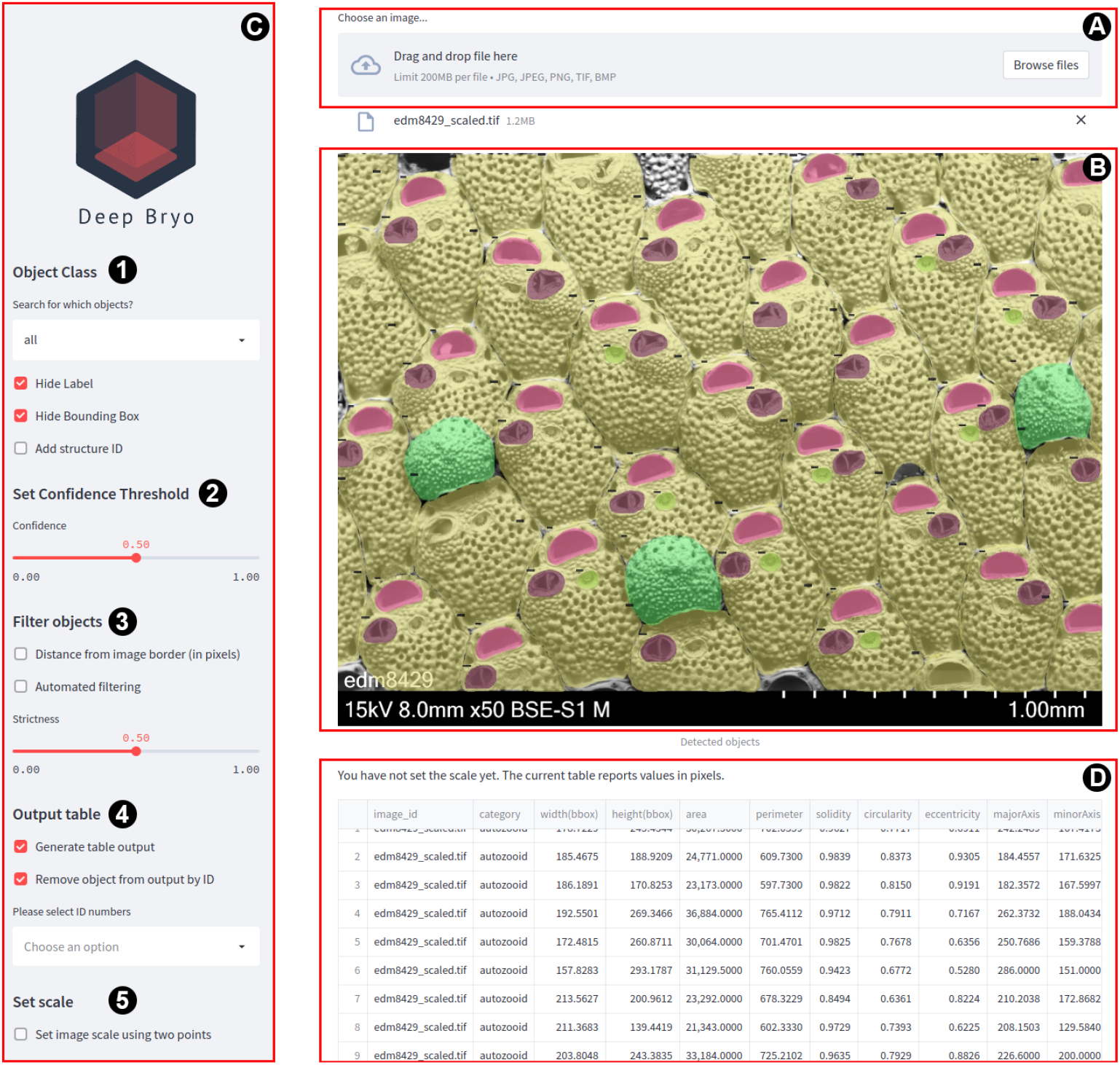
DeepBryo layout as displayed on a web browser. (A) Image upload area; (B) Visualizing the model prediction; (C) Sidebar containing filtering, scaling, and output options; (D) Table output (optional).

**Figure 2.**
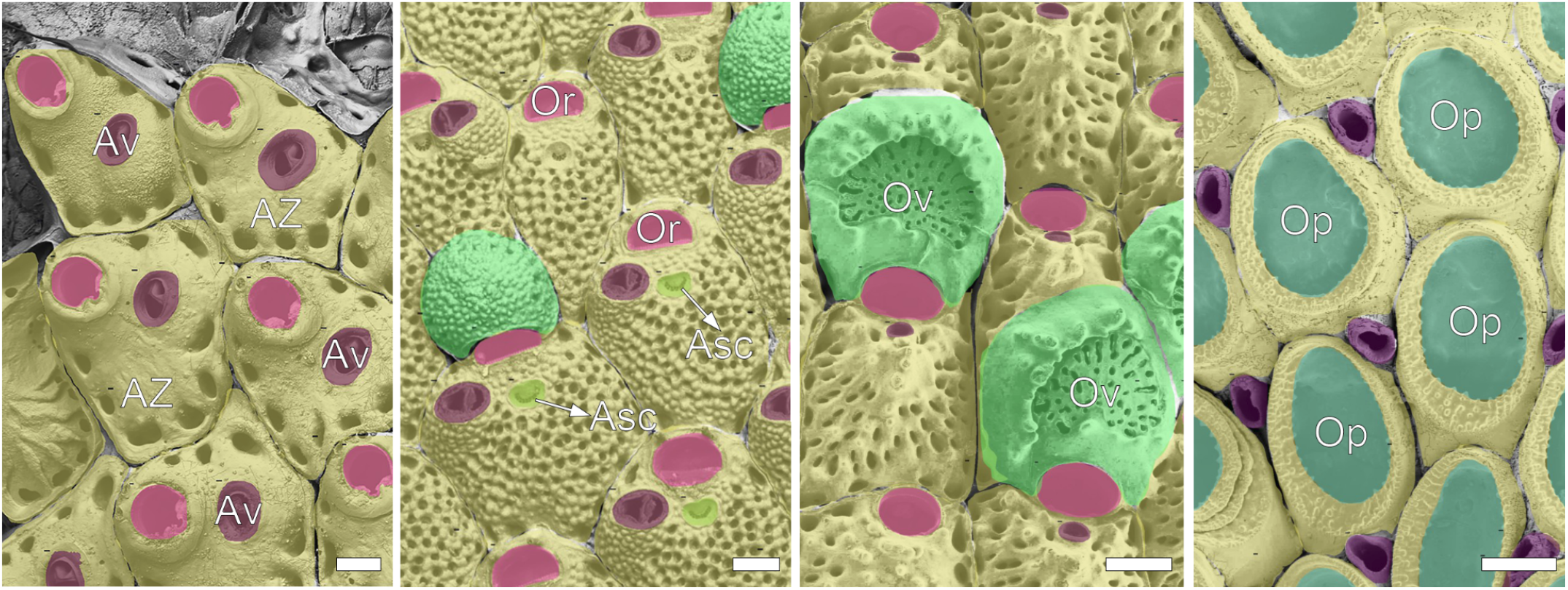
DeepBryo output for unseen images from the validation set. Segmentation masks are overlaid upon the original image using class-specific colors. All six object classes are represented across the four images. AZ (yellow): autozooid; Av (purple): avicularia; Or (pink): orifice; Asc (neon yellow): ascopore; Ov (light green): ovicell; Op (dark green): opesia. See Supplemental File 1 for a glossary of terms. All scale bars are 100 μm.

On the backend, DeepBryo performs instance segmentation and shape measurement as separate tasks. Given an input image, instance segmentation is performed using a Swin Transformer-based Mask R-CNN (Liu et al., 2021). This Swin Transformer model has been fine-tuned using pre-trained weights (ImageNet-1K) (Liu et al., 2021) and an annotated image dataset composed of 935 cheilostome bryozoan images (Table S1), containing 72,412 structures, and captured within the optimal range of magnification (see ***3. Validation*** for performance details).

DeepBryo then performs shape measurement based on the predicted masks using contour functions in the *opencv 4.6.0* python library (Bradski, 2000). The application currently outputs seven commonly used shape descriptors and seven Hu moments derived from each contour’s central moments (Table 1) (Bradski, 2000).

**Table 1.**
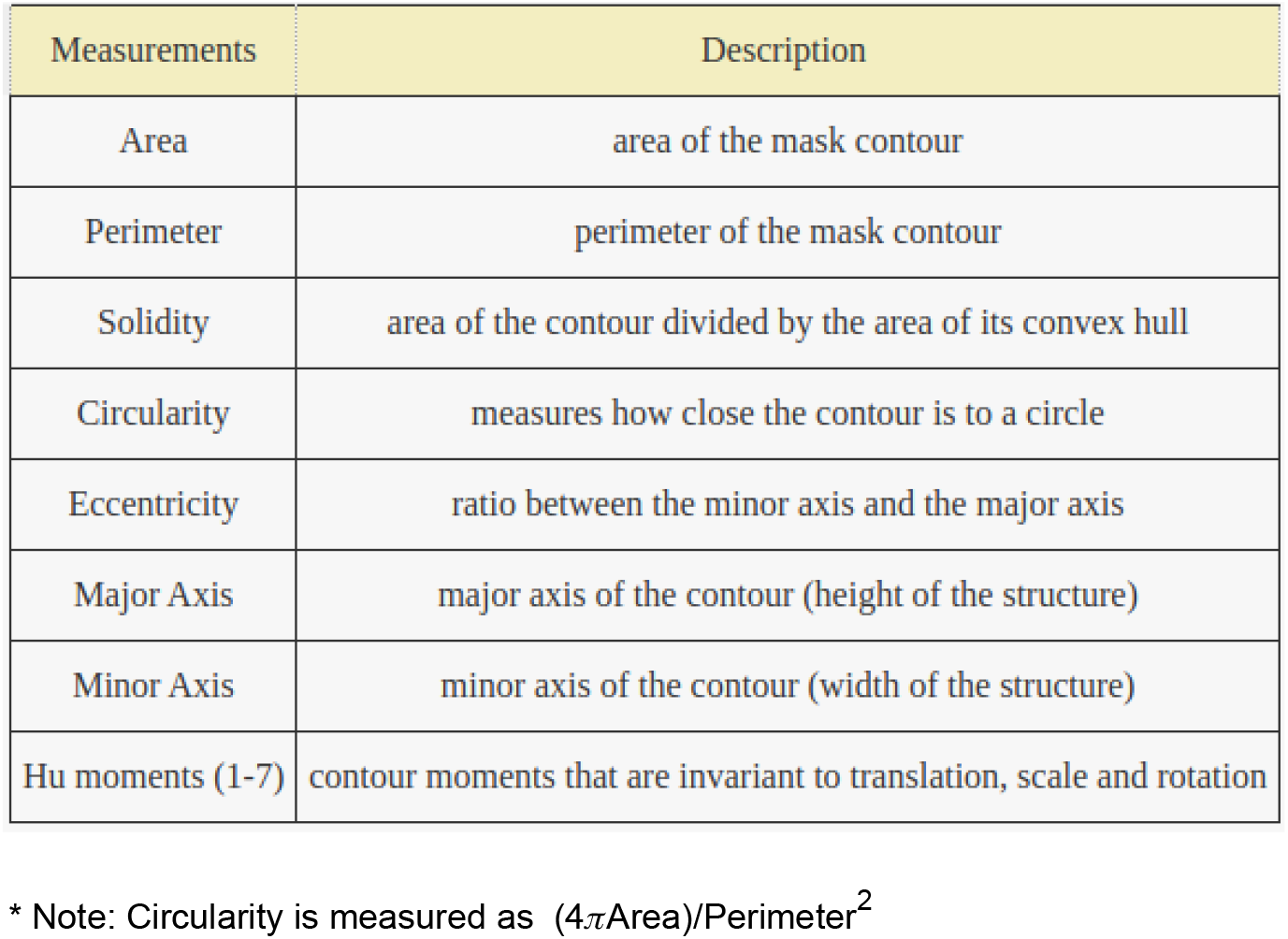
Fourteen contour-based measurements that can be output by the DeepBryo web application for each object. The first seven are basic shape descriptors and the remaining ones are the Hu moments of each contour. Hu Moments, routinely used int the computer vision literature for shape matching, are image moments that are invariant under translations, scale, and rotation.

| Measurements | Description |
| --- | --- |
| Area | area of the mask contour |
| Perimeter | perimeter of the mask contour |
| Solidity | area of the contour divided by the area of its convex hull |
| Circularity | measures how close the contour is to a circle |
| Eccentricity | ratio between the minor axis and the major axis |
| Major Axis | major axis of the contour (height of the structure) |
| Minor Axis | minor axis of the contour (width of the structure) |
| Hu moments (1-7) | contour moments that are invariant to translation, scale and rotation |

\* Note: Circularity is measured as $(4\pi \text{Area}) / \text{Perimeter}^2$

### 2.2 Usage

We here present the steps necessary for using DeepBryo as a web application (https://www.deepbryo.ngrok.io). For command-line interface (CLI) usage, please refer to the official GitHub repository (www.github.com/agporto/DeepBryo).

- Click the “Browse files” button on the right uppermost corner of the screen to upload your image, or simply drag and drop the image into the gray rectangle area next to it (Figure 1A). An upload progress bar will detail how much of the file has been uploaded.
- Once the image is uploaded, the model is automatically initialized and model predictions are shown on the screen as colored masks (Figure 1B). Model predictions have been cached at this point in time, so further steps will not require the model to be reinitialized.
- On the sidebar (Figure 1C), you will find five main sections. In the uppermost section (***Object Class***, Figure 1C-1), you can choose whether to isolate certain object classes using a drop-down menu (“Search for which objects?”). By selecting a specific class, the image masks will be replaced by those belonging exclusively to the class in question. In the same section, you can choose whether to display the bounding boxes corresponding to each detected object. Similarly, you can add a class label or individual structure identification numbers by clicking on checkboxes. By adding the class label, you will also see the model’s confidence in each object’s prediction.
- In the second section (***Set Confidence Threshold***, Figure 1C-2), you can choose whether to filter out objects based on model confidence. The default value is set to 0.5, but can be adjusted based on your filtering needs. Most users will not need to adjust this parameter. Setting this value too low tends to increase the number of spurious detections while setting it too high leads to poor recall performance.
- In the third section (***Filter Objects***, Figure 1C-3), additional filtering options can be enabled using checkboxes. Two main filters are currently available. The “Distance from image border” checkbox filters objects that touch the border of the image. The main purpose of this checkbox is to remove partially incomplete objects from the output. Alternatively, an automated filtering model can further clean the predictions. Automated filtering utilizes a gradient boosting algorithm (Chen & Guestrin, 2016) to classify predictions into “good” or “bad”, “bad” meaning that the object is partially incomplete, not properly cleaned, or simply poorly preserved to be of use. The parameter “Strictness” regulates how strict the gradient boosting algorithm is when filtering out model predictions. Low strictness leads to more filtering out of predictions, while high strictness leads to less filtering out.
- In the fourth section (***Output Table***, Figure 1C-4), you can generate a table output (Figure 1D) containing fourteen traditionally used shape measurements for each detected object using a checkbox. You can also opt to remove certain objects from the table output based on their identification number. To do so, simply type the number in question under “Please select ID numbers” followed by the ‘Enter’ key. Alternatively, select identification numbers using the drop-down menu. Note, however, that the table output will be in pixel units unless you use the next section.
- In the final sidebar section (***Set Scale***, Figure 1C-5), you can scale your measurements using scale bar annotation. While some measurements in DeepBryo are unitless (e.g., eccentricity), others are not (e.g., area, perimeter). If the scale is not set, DeepBryo will output measurements in pixel units. If set, it will output them in millimeters. To set the scale, check ‘Set image scale using two points’ and manually type the scale bar length in μm under ‘Length(μm)’. This will convert the image into a canvas capable of user interaction. You can now place points on the image corresponding to the position and size of the scale bar. You should place one point at the leftmost extreme of the scale bar and another at the rightmost extreme. If the placement is correct, press the ‘Send to streamlit’ button at the left bottom corner of the image. If it is not, press the trash can icon (‘Reset canvas and history’) to have the option of placing new points. Once points have been placed on the scale bar, your table output will be automatically converted to millimeters.

## 3 VALIDATION

### 3.1 Dataset

The dataset used to train and validate the instance segmentation model behind DeepBryo consists of 72,412 structures which were manually annotated on 935 SEM images of bryozoan colonies belonging to ~75% of the families (N=109) and ~30 % of the genera (N=206) within Cheilostomatida (Table S1). The species richness in the dataset is best represented as a range since some taxa could not be identified at the species level from the images available. We estimate that species richness in this dataset ranges from a minimum of ~ 8% (N=443) to a maximum of 9.2% (N=505) of the species diversity among cheilostomes (Table S1). These data have been acquired independently by the coauthors of this manuscript over the years or obtained from public image repositories. They encompass a large array of image quality, resolution, taphonomic degradation, magnification, and equipment used. We consider these pictures to represent most encrusting morphotypes within cheilostomes in which recognition of individual skeletal structures is possible, as well as representative of the images being collected for morphometric research in the group. Some erect species with large flat colony areas are also represented.

We annotated all individual structures contained within the images using polygons. These structures belong to six classes (Figure 2), representing different colony life history traits, such as reproduction, defense, and growth (see Supplemental File 1). Broken, occluded and taphonomically degraded structures were marked as “poor” quality and further used to train a binary automated filtering algorithm based on gradient boosting (Chen & Guestrin, 2016). When training each model, we randomly sampled 90% of the images for training and 10% for validation. The instance segmentation model was trained for 120 epochs, using a learning rate of 10^-4^ and the AdamW optimizer, following general guidelines presented in the original Swin Transformer manuscript (Liu et al., 2021). Image augmentations were performed during training using the *albumentations* library (Buslaev et al., 2020) to increase the model’s generalizability.

### 3.2 Performance

Instance segmentation models can be evaluated in terms of their three underlying tasks: detection, classification, and segmentation. In other words, we can evaluate them in terms of their ability to detect objects (structures) in images, correctly classify them, and produce segmentation masks that have similar morphometric properties to those manually annotated.

#### 3.2.1 Detection

The performance of DeepBryo in terms of object detection can be evaluated using average precision (AP). AP is a common metric used to evaluate object detection performance (Lin et al., 2014). Essentially, it measures the degree to which objects present in the ground-truth data are recovered by the model as valid detections. AP values range from 0 (no recall and/or no precision) to 1 (100% recall and precision). Detections are considered valid if they overlap substantially with manually annotated (ground-truth) objects. As commonly done in the field of computer vision, we use an intersection-over-union value of 0.5 as our threshold for recognizing an object as valid.

In Figure 3, we break down AP values by object class (autozooid, ovicell, etc.) and by object size. We define size according to the area occupied by the object in each image: small (x < 32×32 pixels), medium (32×32 < x < 96×96 pixels), and large (x > 96×96 pixels).

**Figure 3.**
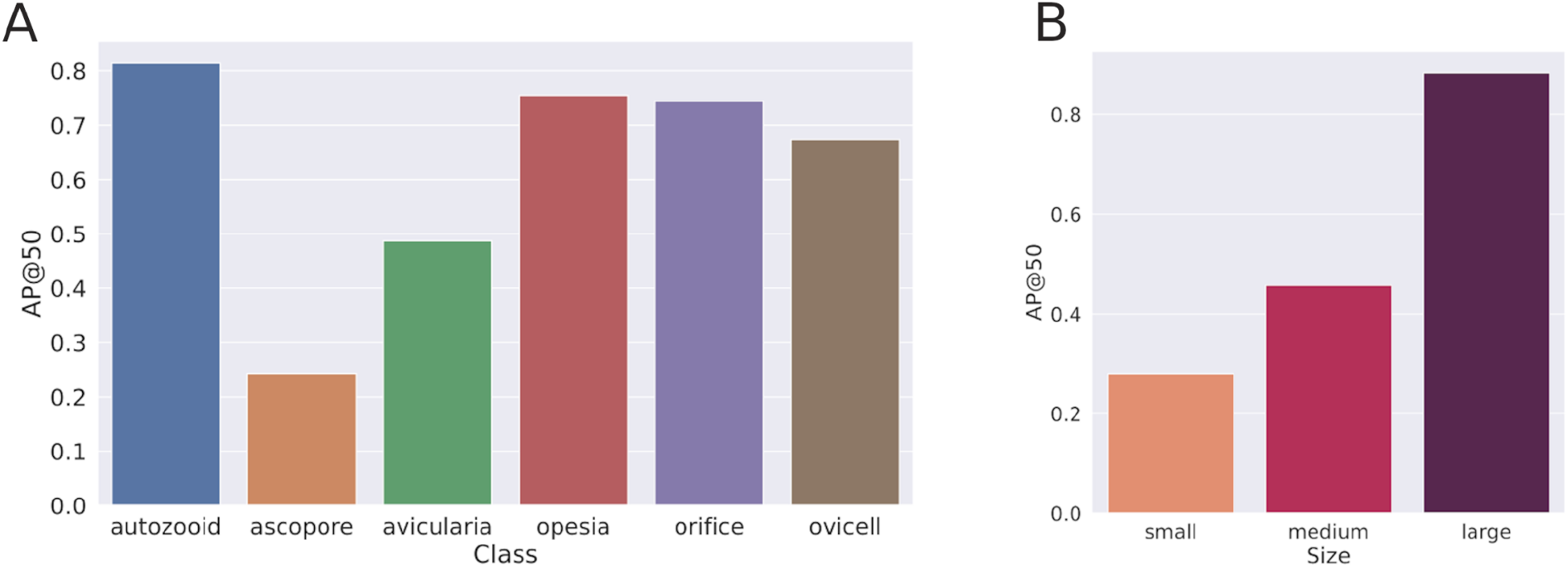
Average precision (AP) of the segmentation masks for an intersection-over-union (IoU) threshold of 50%. (A) AP per object class; (B) AP per object size (small, medium, large). Note the clear association between AP values and object size. Refer to the main text for a description of the size classes and see Supplemental File 1 for a glossary of terms.

As can readily be seen (Figure 3A), AP values are generally high (> 0.7) for most object classes, and decrease considerably with decreased object area (Figure 3B). Autozooids had the highest AP (0.814) among all object classes, followed by opesiae (0.755) and orifices (0.744). This is not surprising, given these are among the largest and most common object types in images of this group. Small structures, such as ascopores, tend to present low levels of AP (0.242) and are, therefore, severely undersampled from images. In other words, the model generally shows high levels of recall and precision as long as objects are large, and vice versa. The same pattern can be observed in the recall and precision curves (Figure S2), which indicate that performance reduction usually occurs through decreased recall rates.

#### 3.2.2 Classification

Once an object is detected, we can measure the extent to which it is correctly classified. Figure 4 illustrates a confusion matrix of object classes. Confusion matrices are a common visualization tool to illustrate the classification performance of machine learning models (Stehman, 1997). It is usually presented as a matrix in which each row represents the percent instances of a ground-truth class that get predicted as belonging to the different classes (columns).

**Figure 4.**
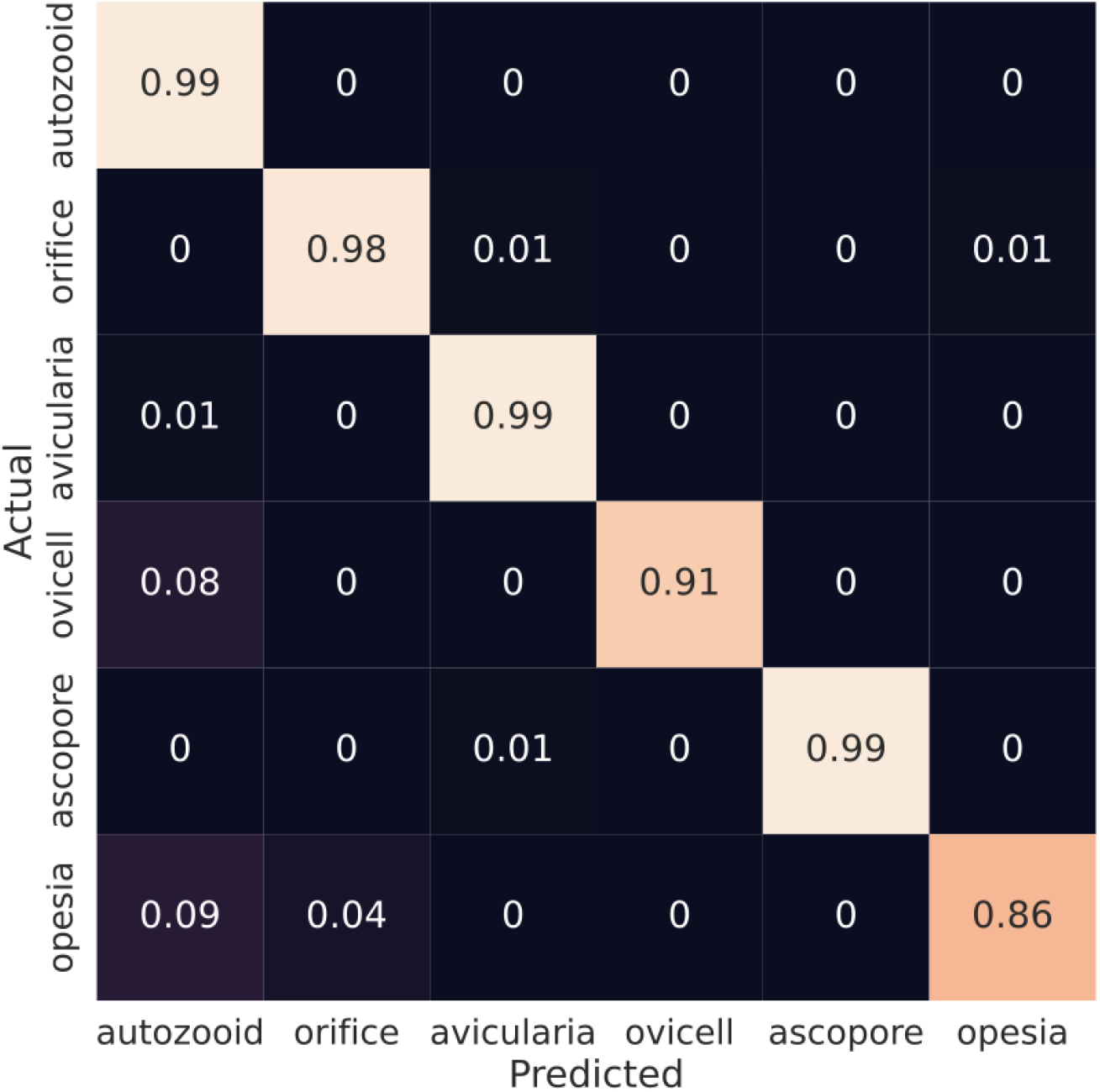
Confusion matrix illustrating the percentage of object predictions falling within each ground-truth object class. Note that the vast majority of object classes are correctly classified by the model.

With the exception of the opesia, class confusion is rather rare and most objects get correctly classified by the model (>0.95). Among opesia objects, a notable percentage (8%) is incorrectly classified as an autozooid. This observation is not surprising given that, in many species, as much as 80-90% of the area of the zooid is occupied by the opesia (see Figure 2). Overall, the observed patterns suggest that classification errors are not a significant source of concern, due to their rarity.

#### 3.2.3 Segmentation

Ultimately, a user that might be interested in DeepBryo will want to know the extent to which the model recovers measurements that are directly comparable to measurements currently used in the literature. To compare the DeepBryo segmentation results with conventional methods, we regressed the predicted area, height (major axis) and width (minor axis) of each detected object against their ground-truth counterparts (Figure 5). We then compare the ability of the model to recover ground-truth measurements based on the coefficient of determination (r^2^) and the relative root-mean-squared error (RMSE) metrics. Finally, we test for deviations from a regression slope of one using the confidence interval of regression of slopes. Deviations from a slope of one would indicate that the model either overestimates (>1) or underestimates (<1) object-level measurements.

**Figure 5.**
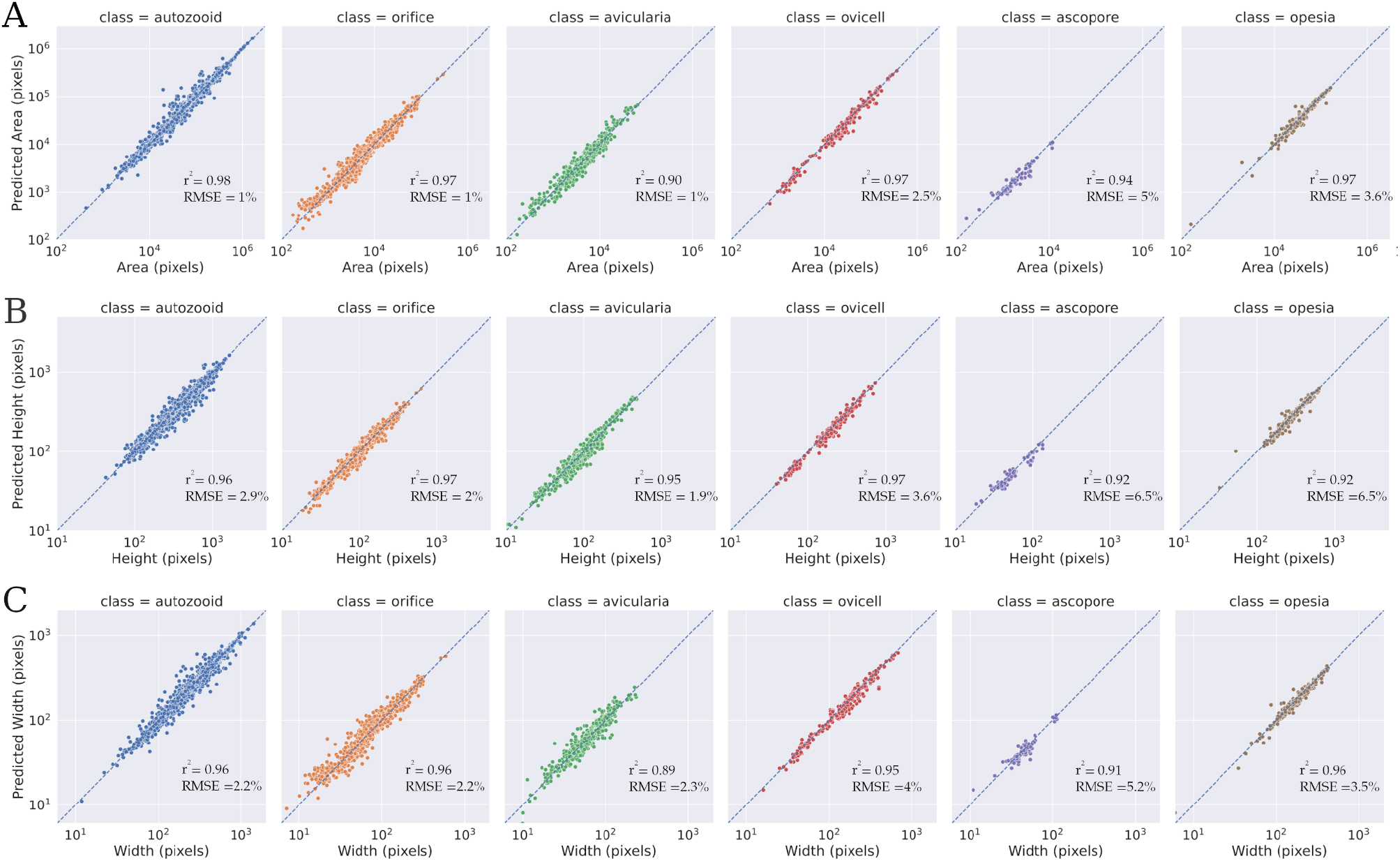
Regression of predicted measurements against their ground truth counterparts. r2 and relative RMSE values are also reported within their respective plot. Rows: (A) Area; (B) Height (contour major axis); (C) Width (contour minor axis).

As seen in Figure 5, all object classes and measurement types have high r^2^ values (>0.89) and low relative RMSE (< or equal to 5% of the range of variation across specimens). Not surprisingly, the confidence interval of the regression slope includes the value of one for all classes, indicating a failure to reject the null hypothesis of a 1:1 relationship between observed and predicted area measurements. Likewise, univariate ANOVAs comparing the means of the distributions of area values per class are equally not significant using p=0.05. In other words, measurements produced by DeepBryo are directly comparable to those obtained manually.

#### 3.2.4 Time

Given the number and diversity of structures, cheilostome bryozoan images take a significant amount of time to be manually annotated. Not surprisingly, taxonomic studies in the group will often collect no more than 20 measurements of each structure, which is considered a fair compromise between the accuracy of the morphometric characterization and the effort in person-hours necessary to do so. Annotation of the entire validation set used in this study took an estimated total of approximately 270 person-hours to complete. In its batch-processing form, as described in the GitHub repository, the model is capable of annotating the same image set in less than a minute. In the web application form, a user can annotate the same set in anywhere between 30 min to 2 person-hours. In other words, DeepBryo reduces the person-hours required for measuring cheilostome bryozoan colonies to less than one percent of the time required to manually annotate such image set.

## 4 DISCUSSION

DeepBryo is a web application for AI-assisted segmentation of SEM images of cheilostome bryozoan colonies (Figure 1). We here demonstrated that it greatly increases the scale and reproducibility of morphometric characterization of bryozoans, as typically done in macroevolutionary, ecological, paleobiological, and taxonomic research. DeepBryo shows high levels of recall and precision for six different structure classes within images (Figure 3) and calculates measurements that are directly comparable to those obtained via expert-based image annotation for a large array of cheilostome genera (Figure 5). Class confusion is rare and is highly concentrated in a single object class (opesia) (Figure 4). Finally, the model has been trained and performs well on a wide array of species of cheilostomes (Table S1).

DeepBryo makes two major contributions to the use of cheilostome bryozoans as a model system in evolution, ecology, and paleobiology. First, it captures (and provides to users) the specialized domain knowledge that is often required to make use of this emerging model system. In our view, the model’s ability to capture domain knowledge will greatly expand the pool of researchers that will be interested in using bryozoans to address important questions in the macroevolutionary, ecological, and paleobiological literature. Second, DeepBryo removes important analytical bottlenecks from morphometric research studies using this clade. For example, a taxonomist interested in capturing the morphological variability within a taxon will be able to do so without making compromises in terms of the number of structures being measured or in terms of the person-hour requirements. Such a taxonomist will also be able to present measurements that can be easily replicated in another laboratory. Similarly, an evolutionary biologist interested in multivariate diversification patterns will be able to do so in a high-throughput and replicable fashion, even if asking questions that require characterizing a wide array of structures and/or species. With small changes in the source code, one could even combine the object detection abilities of DeepBryo with other approaches, such as automated landmarking algorithms (Porto & Voje, 2020). Note, also, that our repository contains all key elements (i.e., building blocks) necessary to adapt the tool to other study systems. The instance segmentation model, for example, can be retrained using a standard COCO dataset (Lin et al., 2014) with simple changes in the configuration file (e.g., change the name of object classes), as described in our GitHub repository.

We do not intend to imply, however, that there are no shortcomings with the current implementation. Most notably, DeepBryo currently focuses on Cheilostomatida. While cheilostomes are the most diverse order within Bryozoa (Bock & Gordon, 2013; Taylor, 2020), containing around 80% of extant species, other orders are also mineralized and could be included in future model retraining. Therefore, future work will concentrate on increasing taxonomic coverage and expanding the app to other bryozoan orders. Likewise, the object detection performance of DeepBryo has clear room for improvement. Currently, it focuses on images obtained within the optimal magnification range for morphometric analyses (between 30x and 90x magnification) and therefore shows poor recall for small objects. It could, however, be adapted to lower magnification levels through the use of a sliding window approach, in which one large image gets subdivided into smaller high-resolution sub-windows. We generally consider larger magnifications (>90x) as unlikely to be employed in morphometric studies of whole colonies, as the entire image would only encompass partial structures. Another significant challenge relates to the scale bars. Currently, DeepBryo utilizes manual digitization of scale bars. Automated recognition of scale bar size would reduce the need for human intervention during the prediction process. Finally, Deepbryo cannot account for discrepancies in the standards of different fields. For example, in macroevolutionary work, researchers tend to include the cystid as part of avicularia-related measurements. In taxonomic work, researchers will often keep the cystid as a separate entity and measure only the mandible and the proximal membranous part. Currently, DeepBryo adopts the standard used in macroevolutionary studies.

In summary, DeepBryo is a new web application that enables rapid morphometric characterization of cheilostome bryozoan images captured using SEM machines. The application has a simple web interface that requires no pre-processing of the image data. It enables users to predict, visualize, filter, and download segmentation results. With further image annotation and user feedback, we expect DeepBryo to grow significantly over time, providing researchers from different areas of biology the ability to use this emerging model system in tackling important research questions in their respective fields.

## AUTHORS’ CONTRIBUTIONS

AP and EDM conceived the study; AP trained the model and developed the application with inputs from all authors; All authors captured images, tested the application, and defined the taxonomy; AP, HLR, MHR and EDM annotated images; AP and EDM wrote the manuscript. All authors contributed to the manuscript revision.

## ACKNOWLEDGEMENTS

This work was supported in part by grants from the Research Council of Norway (grant 314499 to EDM) and from NVIDIA Corporation (Nvidia Hardware Grant to AP). LHL and MHR received funding from the European Research Council under the European Union’s Horizon 2020 research and innovation program (grant agreement no. 724324 to LHL).

## DATA AVAILABILITY STATEMENT

The entire training/validation dataset will be made available in appropriate repositories upon publication. A live server is available at https://www.deepbryo.ngrok.io. The DeepBryo source code can be found at https://www.github.com/agporto/DeepBryo.

