## Supplemental Material for "DeepBryo: a web app for AI-assisted morphometric characterization of cheilostome bryozoans"

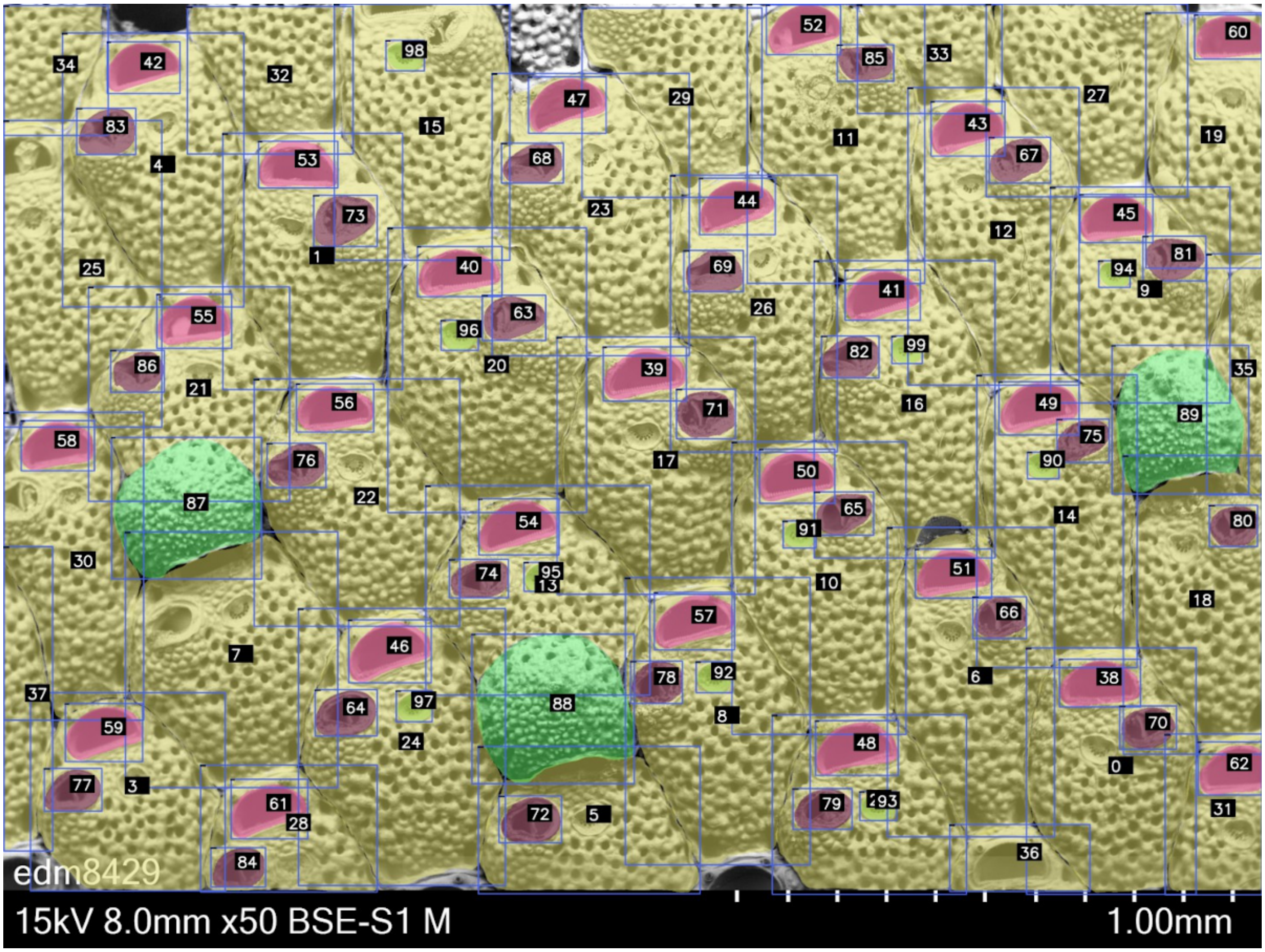


Figure S1 - Prediction layout shown by the DeepBryo app when activating the option to show bounding boxes (blue) and the structure-level identification numbers (squares with dark background). ID numbers can be used to filter the output table.

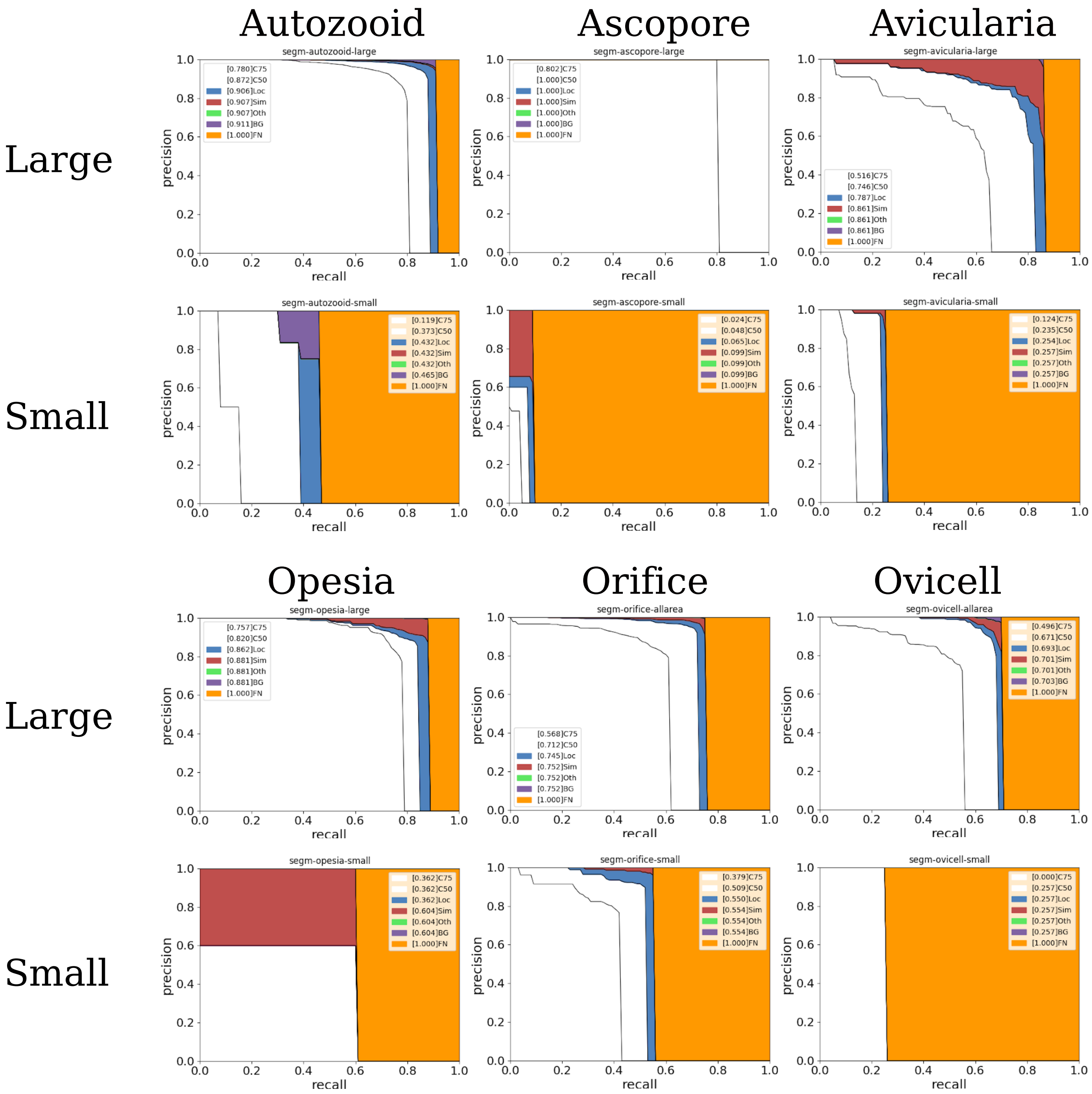


Figure S2 - Precision-recall (PR) curves for all six classes of objects, broken down according to structure size (small vs large). The area in white defines the PR curve for an IoU value of 0.5. By adding the blue area (Loc), it becomes the PR curve while ignoring localization error (IoU of 0.1). By adding the area in red (Sim), it becomes the PR curve while ignoring classification errors (e.g. mistaking an ovicell as an autozooid).By adding the area in purple (BG), it becomes the PR curve including false positive errors. Finally, the area in orange (FN) represents the false negatives.

### Glossary

*Autozooid*: essential unit of a bryozoan colony. Equipped with a polypide devoted to feeding and other somatic functions.

*Avicularium*: a specialized zooid with a reduced, non-feeding polypide and a mandible with mechano-sensory capabilities. Involved in the defense against predators and cleaning of the colony.

*Ascopore*: frontal pore which operates as an inlet to the ascus, a flexible sac pertaining to the plumbing system of ascophoran-grade cheilostome bryozoans.

*Encrusting*: growth mode in which the colony expands like a sheet adpressed to the surface of the substratum.

*Kenozooid*: a specialized zooid that lacks a polypide. Involved in many structural functions, such as space-filling, attachment, and colony support.

*Opesia*: opening below the frontal membrane in anascan-grade cheilostome bryozoans lacking a calcified frontal shield.

*Polymorphism*: variation in zooid morphology reflecting division of labor within the colony.

*Orifice*: opening in the frontal shield of autozooids through which the lophophore can be extruded.

*Ovicell*: brood chamber found in cheilostomes.

**Table S1 – List of Bryozoa Cheilostomatida taxa included in the dataset used to train/validate the instance segmentation model.**

| Taxon_name | Authorship | Genus | Worms::Family |
|---|---|---|---|
| *Acanthodesia* sp. | | *Acanthodesia* | Membraniporidae |
| *Actisecos regularis* | Canu & Bassler, 1927 | *Actisecos* | Actisecidae |
| *Actisecos* sp. 1 | sensu Di Martino & Taylor, 2015 | *Actisecos* | Actisecidae |
| *Adelascopora secunda* | Hayward & Thorpe, 1988 | *Adelascopora* | Fenestrulinidae |
| *Adeona* sp. | | *Adeona* | Adeonidae |
| *Adeonella haywardii* | Amui, 2005 | *Adeonella* | Adeonidae |
| *Adeonella intricaria* | Busk, 1884 | *Adeonella* | Adeonidae |
| *Adeonella lichenoides* | (Lamarck, 1816) | *Adeonella* | Adeonidae |
| *Adeonella pallasii* | (Heller, 1867) | *Adeonella* | Adeonidae |
| *Adeonella polystomella* | (Reuss, 1847) | *Adeonella* | Adeonidae |
| *Adeonella* sp. | | *Adeonella* | Adeonidae |
| *Adeonellopsis* cf. *distoma* | (Busk, 1858) | *Adeonellopsis* | Adeonidae |
| *Adeonellopsis foliacea* | MacGillivray, 1886 | *Adeonellopsis* | Adeonidae |
| *Adeonellopsis japonica* | (Ortmann, 1890) | *Adeonellopsis* | Adeonidae |
| *Adeonellopsis pentapora* | Canu & Bassler, 1929 | *Adeonellopsis* | Adeonidae |
| *Adeonellopsis* spp. | | *Adeonellopsis* | Adeonidae |
| *Aechmella* sp. | | *Aechmella* | Onychocellidae |
| *Aimulosia marsupium* | (MacGillivray, 1869) | *Aimulosia* | Buffonellodidae |
| *Aimulosia* sp. | | *Aimulosia* | Buffonellodidae |
| *Akatopora* cf. *leucocypha* | (Marcus, 1937) | *Akatopora* | Antroporidae |
| *Alderina imbellis* | (Hincks, 1860) | *Alderina* | Calloporidae |
| *Amphiblestrum auritum* | (Hincks, 1877) | *Amphiblestrum* | Calloporidae |
| *Amphiblestrum contentum* | Gordon, 1986 | *Amphiblestrum* | Calloporidae |
| *Amphiblestrum trifolium* | (Wood, 1844) | *Amphiblestrum* | Calloporidae |
| *Anaskopora* sp. | | *Anaskopora* | Cribrilinidae |
| *Anchicleidochasma mirabile* | (Harmer, 1957) | *Anchicleidochasma* | Cleidochasmatidae |
| *Antarcticaetos bubeccata* | (Rogick, 1955) | *Antarcticaetos* | Romancheinidae |
| *Antarctothoa antarctica* | Moyano & Gordon, 1980 | *Antarctothoa* | Hippothoidae |
| *Antarctothoa haywardi* | Kuklinski & Barnes, 2009 | *Antarctothoa* | Hippothoidae |
| *Antoniettella exigua* | Di Martino & Taylor, 2012 | *Antoniettella* | Cribrilinidae |
| *Antropora* cf. *granulifera* | (Hincks, 1880) | *Antropora* | Antroporidae |
| *Antropora* cf. *minor* | (Hincks, 1880) | *Antropora* | Antroporidae |
| *Antropora granulifera* | (Hincks, 1880) | *Antropora* | Antroporidae |
| *Antropora minor* | (Hincks, 1880) | *Antropora* | Antroporidae |
| *Antropora* cf. *subvespertilio* | (Canu & Bassler, 1929) | *Antropora* | Antroporidae |
| *Antropora typica* | Canu & Bassler, 1928 | *Antropora* | Antroporidae |
| *Arachnopusia columnaris* | Hayward & Thorpe, 1988 | *Arachnopusia* | Arachnopusiidae |
| *Arachnopusia perforata* | (Maplestone, 1909) | *Arachnopusia* | Arachnopusiidae |
| *Arachnopusia quadralabia* | Uttley & Bullivant, 1972 | *Arachnopusia* | Arachnopusiidae |
| *Arachnopusia* sp. | | *Arachnopusia* | Arachnopusiidae |
| *Arthropoma* n. sp. | | *Arthropoma* | Lacernidae |
| *Arthropoma* sp. 1 | | Inapplicable | Inapplicable |
| Ascophora indet. 1 | | Inapplicable | Inapplicable |
| Ascophora indet. 2 | | Inapplicable | Inapplicable |
| Ascophora indet. 3 | | Inapplicable | Inapplicable |
| Ascophora indet. 4 | | Inapplicable | Inapplicable |
| *Atlantisina acantha* | Berning, Harmelin & Bader, 2017 | *Atlantisina* | Atlantisinidae |
| *Atlantisina atlantis* | Berning, Harmelin & Bader, 2017 | *Atlantisina* | Atlantisinidae |
| *Atlantisina gorringensis* | Berning, Harmelin & Bader, 2017 | *Atlantisina* | Atlantisinidae |
| *Atlantisina lionensis* | Berning, Harmelin & Bader, 2017 | *Atlantisina* | Atlantisinidae |
| *Atlantisina meteor* | Berning, Harmelin & Bader, 2017 | *Atlantisina* | Atlantisinidae |
| *Atlantisina seinensis* | Berning, Harmelin & Bader, 2017 | *Atlantisina* | Atlantisinidae |
| *Atlantisina tricornis* | Berning, Harmelin & Bader, 2017 | *Atlantisina* | Atlantisinidae |
| *Bathycyclopora suroiti* | Berning, Harmelin & Bader, 2017 | *Bathycyclopora* | Atlantisinidae |
| *Bathycyclopora vibraculata* | (Calvet, 1931) | *Bathycyclopora* | Atlantisinidae |
| *Beania* cf. *plurispinosa* | Uttley & Bullivant, 1972 | *Beania* | Beaniidae |
| *Beania elongata* | (Hincks, 1885) | *Beania* | Beaniidae |
| *Biflustra* cf. *savartii* | (Audouin, 1826) | *Biflustra* | Membraniporidae |
| *Biflustra limosa* | (Waters, 1909) | *Biflustra* | Membraniporidae |
| *Biflustra denticulata* | (Busk, 1856) | *Biflustra* | Membraniporidae |
| *Biflustra* n. sp. | | *Biflustra* | Membraniporidae |
| *Biflustra savartii* | (Audouin, 1826) | *Biflustra* | Membraniporidae |
| *Bitectipora rostrata* | (MacGillivray, 1887) | *Bitectipora* | Bitectiporidae |
| *Bracebridgia* sp. | | *Bracebridgia* | Adeonidae |
| *Bragella pseudofedora* | Di Martino, Taylor, Cotton & Pearson, 2017 | *Bragella* | Ascosiidae |
| *Bryopesanser baderae* | Tilbrook, 2012 | *Bryopesanser* | Escharinidae |
| *Bryopesanser* cf. *serratus* | Dick, Tilbrook & Mawatari, 2006 | *Bryopesanser* | Escharinidae |
| *Bryopesanser pesanseris* | (Smitt, 1873) | *Bryopesanser* | Escharinidae |
| *Bryopesanser sanfilippoae* | Di Martino & Taylor, 2015 | *Bryopesanser* | Escharinidae |
| *Buffonellaria acorensis* | Berning & Kukliński, 2008 | *Buffonellaria* | Celleporidae |

| | | | |
|---|---|---|---|
| *Buffonellaria antoniettae* | Berning & Kukliński, 2008 | *Buffonellaria* | Celleporidae |
| *Buffonellaria biaperta* | sensu Hayami, 1975 | *Buffonellaria* | Celleporidae |
| *Buffonellaria harmelini* | Berning & Kukliński, 2008 | *Buffonellaria* | Celleporidae |
| *Buffonellaria muriella* | Berning & Kukliński, 2008 | *Buffonellaria* | Celleporidae |
| *Buffonellaria turbula* | Gordon, 1989 | *Buffonellaria* | Celleporidae |
| *Callopora horridoidea* | Androsova, 1958 | *Callopora* | Calloporidae |
| Calloporidae spp. | | Inapplicable | Calloporidae |
| *Calloporina angustipora* | (Hincks, 1885) | *Calloporina* | Microporellidae |
| *Calloporina decorata* | (Reuss, 1848) | *Calloporina* | Microporellidae |
| *Calloporina* sp. | | *Calloporina* | Microporellidae |
| *Calvetopora inflata* | (Calvet, 1906) | *Calvetopora* | Atlantisinidae |
| *Calvetopora otapostasis* | Berning, Harmelin & Bader, 2017 | *Calvetopora* | Atlantisinidae |
| *Calyptotheca anceps* | (MacGillivray, 1879) | *Calyptotheca* | Lanceoporidae |
| *Calyptotheca* cf. *inclusa* | (Thornely, 1906) | *Calyptotheca* | Lanceoporidae |
| *Calyptotheca hastingsae* | Harmer, 1957 | *Calyptotheca* | Lanceoporidae |
| *Calyptotheca rugosa* | Hayward, 1974 | *Calyptotheca* | Lanceoporidae |
| *Calyptotheca* spp. | | *Calyptotheca* | Lanceoporidae |
| *Cellaria harmelini* | d'Hondt, 1973 | *Cellaria* | Cellariidae |
| *Cellaria immersa* | (Tenison Woods, 1880) | *Cellaria* | Cellariidae |
| *Cellaria melillensis* | el-Hajjaji, 1987 | *Cellaria* | Cellariidae |
| *Calyptotheca* cf. *nigra* | Dumont, 1981 | Inapplicable | Inapplicable |
| *Cellaria* sp. | | *Cellaria* | Cellariidae |
| *Cellarinella nutti* | Rogick, 1956 | *Cellarinella* | Sclerodomidae |
| *Cellarinella* sp. | | *Cellarinella* | Sclerodomidae |
| *Celleporaria aperta* | (Hincks, 1882) | *Celleporaria* | Lepraliellidae |
| *Celleporaria* cf. *vermiformis* | (Waters, 1909) | *Celleporaria* | Lepraliellidae |
| *Celleporaria* n. sp. | | *Celleporaria* | Lepraliellidae |
| *Celleporaria* sp. | | *Celleporaria* | Lepraliellidae |
| *Celleporaria vermiformis* | (Waters, 1909) | *Celleporaria* | Lepraliellidae |
| *Celleporella hyalina* | (Linnaeus, 1767) | *Celleporella* | Hippothoidae |
| *Celleporella* sp. | | *Celleporella* | Hippothoidae |
| *Celleporina* cf. *costazii* | (Audouin, 1826) | *Celleporina* | Celleporidae |
| *Celleporina* cf. *tropica* | Hayward, 1988 | *Celleporina* | Celleporidae |
| *Celleporina grandis* | Gordon, 1989 | *Celleporina* | Celleporidae |
| *Celleporina* sp. | | *Celleporina* | Celleporidae |
| *Chaperia granulosa* | Gordon, 1986 | *Chaperia* | Chaperiidae |
| *Chaperia infundibulata* | d'Hondt, 1988 | *Chaperia* | Chaperiidae |
| *Chaperia* sp. | | *Chaperia* | Chaperiidae |
| *Chaperiopsis* n. sp. | | *Chaperiopsis* | Chaperiidae |
| *Chaperiopsis rubida* | (Hincks, 1881) | *Chaperiopsis* | Chaperiidae |
| *Chaperiopsis* sp. | | *Chaperiopsis* | Chaperiidae |
| *Chaperiopsis spiculata* | Uttley, 1949 | *Chaperiopsis* | Chaperiidae |
| *Characodoma granosum* | (Seguenza, 1880) | *Characodoma* | Cleidochasmatidae |
| *Characodoma longitudinale* | (Harmer, 1957) | *Characodoma* | Cleidochasmatidae |
| *Characodoma* spp. | | *Characodoma* | Cleidochasmatidae |
| *Cheiloporina clarksvillensis* | Di Martino, Taylor & Portell, 2017 | *Cheiloporina* | Cheiloporinidae |
| *Chiastosella enigma* | Brown, 1954 | *Chiastosella* | Escharinidae |
| *Chiastosella* n. sp. | | *Chiastosella* | Escharinidae |
| *Chorizopora brongniarti* | (Audouin, 1826) | *Chorizopora* | Chorizoporidae |
| *Cigclisula occlusa* | (Busk, 1884) | *Cigclisula* | Colatooeciidae |
| *Cigclisula* sp. 2 | sensu Di Martino & Taylor, 2015 | *Cigclisula* | Colatooeciidae |
| *Cleidochasmidra portisi* | (Neviani, 1895) | *Cleidochasmidra* | Cleidochasmatidae |
| *Coleopora* sp. | | *Coleopora* | Teuchoporidae |
| *Conescharellina* sp. | | *Conescharellina* | Conescharellinidae |
| *Conopeum* cf. *lacroixi* | (Audouin, 1826) | *Conopeum* | Electridae |
| *Conopeum lacroixii* | (Audouin, 1826) | *Conopeum* | Electridae |
| *Conopeum nakanosum* | Grischenko, Dick & Mawatari, 2007 | *Conopeum* | Electridae |
| *Conopeum seurati* | (Canu, 1928) | *Conopeum* | Electridae |
| *Corbulipora tubulifera* | (Hincks, 1881) | *Corbulipora* | Cribrilinidae |
| *Cosciniopsis* cf. *lonchaea* | (Busk, 1884) | *Cosciniopsis* | Gigantoporidae |
| *Cranosina* cf. *coronata* | (Hincks, 1881) | *Cranosina* | Calloporidae |
| *Cranosina coronata* | (Hincks, 1881) | *Cranosina* | Calloporidae |
| *Crassimarginatella bobiesi* | (David & Pouyet, 1974) | *Crassimarginatella* | Calloporidae |
| *Crassimarginatella cucullata* | (Waters, 1898) | *Crassimarginatella* | Calloporidae |
| *Crassimarginatella* sp. 1 | | *Crassimarginatella* | Calloporidae |
| *Crepidacantha bracebridgei* | Brown, 1954 | *Crepidacantha* | Crepidacanthidae |
| *Crepidacantha poissonii* | (Audouin, 1826) | *Crepidacantha* | Crepidacanthidae |
| *Crepidacantha* sp. | | *Crepidacantha* | Crepidacanthidae |
| *Cribellopora* cf. *latigastra* | (David, 1949) | *Cribellopora* | Lacernidae |
| *Cribellopora* n. sp. | | *Cribellopora* | Lacernidae |
| *Cribellopora* sp. | | *Cribellopora* | Lacernidae |
| *Cribrilina (Cribrilina) cryptooecium* | Norman, 1903 | *Cribrilina* | Cribrilinidae |
| *Cribrilina spitsbergensis* | Norman, 1903 | *Cribrilina* | Cribrilinidae |

| | | | |
|---|---|---|---|
| Cribrilinidae indet. | | Inapplicable | Cribrilinidae |
| *Cucullipora* sp. | | *Cucullipora* | Adeonidae |
| *Cupuladria bugei* | Galopim de Carvalho, 1966 | *Cupuladria* | Cupuladriidae |
| *Cupuladria cavernosa* | Cadée, 1979 | *Cupuladria* | Cupuladriidae |
| *Cupuladria* spp. | | *Cupuladria* | Cupuladriidae |
| *Cyclicopora praelonga* | Hincks, 1884 | *Cyclicopora* | Cyclicoporidae |
| *Cyclocolposa perforata* | Canu & Bassler, 1923 | *Cyclocolposa* | Bryocryptellidae |
| *Cycloperiella rubra* | Canu & Bassler, 1923 | *Cycloperiella* | Stomachetosellidae |
| *Dakariella concinna* | Hayward, 1993 | *Dakariella* | Smittinidae |
| *Discoporella* spp. | | *Discoporella* | Cupuladriidae |
| *Discoporella umbellata* | (Defrance, 1823) | *Discoporella* | Cupuladriidae |
| *Discoradius rutella* | (Tenison Woods, 1880) | *Discoradius* | Lunulitidae |
| *Discoradius tanzaniensis* | Di Martino, Taylor, Cotton & Pearson, 2017 | *Discoradius* | Lunulitidae |
| *Drepanophora* cf. *indica* | Hayward, 1988 | *Drepanophora* | Celleporidae |
| *Drepanophora indica* | Hayward, 1988 | *Drepanophora* | Celleporidae |
| *Ehrhardina voigti* | Martha & Taylor, 2016 | *Ehrhardina* | Onychocellidae |
| *Electra pilosa* | (Linnaeus, 1767) | *Electra* | Electridae |
| *Electra* sp. | | *Electra* | Electridae |
| *Elleschara* cf. *rylandi* | (Soule, Soule & Chaney, 1995) | *Elleschara* | Romancheinidae |
| *Ellisina* aff. *andense* | (Hayami, 1975) | *Ellisina* | Calloporidae |
| *Ellisina sericea* | (MacGillivray, 1890) | *Ellisina* | Calloporidae |
| *Emballotheca longidens* | (Cipolla, 1921) | *Emballotheca* | Lanceoporidae |
| *Eminooecia carsonae* | (Rogick, 1957) | *Eminooecia* | Eminooeciidae |
| *Entomaria spinifera* | Canu, 1914 | *Entomaria* | Aspidostomatidae |
| *Escharella* cf. *klugei* | Hayward, 1979 | *Escharella* | Escharellidae |
| *Escharella longicollis* | (Jullien, 1882) | *Escharella* | Escharellidae |
| *Escharella plana* | (Canu & Bassler, 1920) | *Escharella* | Escharellidae |
| *Escharella reussiana* | (Busk, 1859) | *Escharella* | Escharellidae |
| *Escharella diaphana* | (MacGillivray, 1879) | *Escharella* | Escharellidae |
| *Escharella ventricosa* | (Hassall, 1842) | *Escharella* | Escharellidae |
| *Escharoides coccinea* | (Abildgaard, 1806) | *Escharoides* | Exochellidae |
| *Escharoides costifer* | (Osburn, 1914) | *Escharoides* | Exochellidae |
| *Escharoides longirostris* | Dumont, 1981 | *Escharoides* | Exochellidae |
| *Escharoides mamillata* | (Wood, 1844) | *Escharoides* | Exochellidae |
| *Escharoides* n. sp. | | *Escharoides* | Exochellidae |
| *Eurystomella foraminigera* | (Hincks, 1883) | *Eurystomella* | Eurystomellidae |
| *Euthyroides* n. sp. | | *Euthyroides* | Euthyroididae |
| *Exechonella brasiliensis* | Canu & Bassler, 1928 | *Exechonella* | Exochellidae |
| *Exechonella* cf. *brasiliensis* | Canu & Bassler, 1928 | *Exechonella* | Exochellidae |
| *Exechonella* n. sp. | | *Exechonella* | Exochellidae |
| *Exechonella* spp. | | *Exechonella* | Exochellidae |
| *Exochella armata* | (Hincks, 1882) | *Exochella* | Exochellidae |
| *Exochella hymanae* | (Rogick, 1956) | *Exochella* | Exochellidae |
| *Fedora* aff. *nodosa* | Silén, 1947 | *Fedora* | Ascosiidae |
| *Fenestrulina antarctica* | Hayward & Thorpe, 1990 | *Fenestrulina* | Fenestrulinidae |
| *Fenestrulina candida* | (MacGillivray, 1860) | *Fenestrulina* | Fenestrulinidae |
| *Fenestrulina delicia* | Winston, Hayward & Craig, 2000 | *Fenestrulina* | Fenestrulinidae |
| *Fenestrulina fritilla* | Hayward & Ryland, 1990 | *Fenestrulina* | Fenestrulinidae |
| *Fenestrulina jocunda* | Hayward & Ryland, 1990 | *Fenestrulina* | Fenestrulinidae |
| *Fenestrulina parviporus* | Dick & Grischenko, 2016 | *Fenestrulina* | Fenestrulinidae |
| *Fenestrulina proxima* | (Waters, 1904) | *Fenestrulina* | Fenestrulinidae |
| *Fenestrulina rugula* | Hayward & Ryland, 1990 | *Fenestrulina* | Fenestrulinidae |
| *Fenestrulina* spp. | | *Fenestrulina* | Fenestrulinidae |
| *Figularia carinata* | (Waters, 1887) | *Figularia* | Cribrilinidae |
| *Figularia* cf. *figularis* | (Johnston, 1847) | *Figularia* | Cribrilinidae |
| *Figularia* sp. 1 | | *Figularia* | Cribrilinidae |
| *Flabellopora* sp. | | *Flabellopora* | Conescharellinidae |
| *Floridina multilamellosa* | Voigt, 1924 | *Floridina* | Onychocellidae |
| *Floridina regularis* | Canu & Bassler, 1923 | *Floridina* | Onychocellidae |
| *Fodinella* sp. | | *Fodinella* | Phidoloporidae |
| *Fovoporella spectabilis* | (Hincks, 1891) | *Fovoporella* | Schizoporellidae |
| *Galeopsis polyporus* | (Brown, 1952) | *Galeopsis* | Celleporidae |
| *Galeopsis porcellanicus* | (Hutton, 1873) | *Galeopsis* | Celleporidae |
| *Gemelliporella* n. sp. | | *Gemelliporella* | Gemelliporellidae |
| *Gemelliporidra* sp. | | *Gemelliporidra* | Schizoporellidae |
| *Gigantopora pupa* | (Jullien, 1903) | *Gigantopora* | Gigantoporidae |
| *Hemiphylactella pulchra* | Vigneaux, 1949 | *Hemiphylactella* | Escharellidae |
| *Hemismittina pustulosa* | Vigneaux, 1949 | *Hemismittina* | Bryocryptellidae |
| *Herentia andreasi* | Berning, Tilbrook & Rosso, 2008 | *Herentia* | Escharinidae |
| *Herentia hyndmanni* | (Johnston, 1847) | *Herentia* | Escharinidae |
| *Herentia thalassae* | David & Pouyet, 1978 | *Herentia* | Escharinidae |
| *Hippadenella inerma* | (Calvet, 1909) | *Hippadenella* | Buffonellidae |
| *Hippaliosina acutirostris* | Canu & Bassler, 1929 | *Hippaliosina* | Hippaliosinidae |

| | | | |
|---|---|---|---|
| *Hippaliosina rostrigera* | (Smitt, 1873) | *Hippaliosina* | Hippaliosinidae |
| *Hippaliosina sandbergeri* | (Reuss, 1869) | *Hippaliosina* | Hippaliosinidae |
| *Hippaliosina triforma* | Canu & Bassler, 1929 | *Hippaliosina* | Hippaliosinidae |
| *Hippellozoon novaezelandiae* | (Waters, 1895) | *Hippellozoon* | Phidoloporidae |
| *Hippomenella vellicata* | (Hutton, 1873) | *Hippomenella* | Escharinidae |
| *Hippomenella* spp. | | *Hippomenella* | Romancheinidae |
| *Hippopleurifera sedgwicki* | (Milne Edwards, 1838) | *Hippopleurifera* | Romancheinidae |
| *Hippopodina* cf. *iririkiensis* | Tilbrook, 1999 | *Hippopodina* | Hippopodinidae |
| *Hippopodina iririkiensis* | Tilbrook, 1999 | *Hippopodina* | Hippopodinidae |
| *Hippopodina pulcherrima* | (Canu & Bassler, 1928) | *Hippopodina* | Hippopodinidae |
| *Hippopodina* spp. | | *Hippopodina* | Hippopodinidae |
| *Hippoporella cornuta* | (Busk, 1859) | *Hippoporella* | Hippoporidridae |
| *Hippoporidra edax* | (Busk, 1859) | *Hippoporidra* | Hippoporidridae |
| *Hippothoa* cf. *calciophilia* | Gordon, 1984 | *Hippothoa* | Hippothoidae |
| *Hippotrema janthina* | (Smitt, 1873) | *Hippotrema* | Hippoporidridae |
| *Inversiula nutrix* | Jullien 1888 | *Inversiula* | Inversiulidae |
| *Iodictyum yaldwyni* | Powell, 1967 | *Iodictyum* | Phidoloporidae |
| *Isoschizoporella similis* | Hayward & Thorpe, 1988 | *Isoschizoporella* | Eminooeciidae |
| *Isoschizoporella virgula* | Hayward & Thorpe, 1988 | *Isoschizoporella* | Eminooeciidae |
| *Jellyella eburnea* | (Hincks, 1891) | *Jellyella* | Membraniporidae |
| *Juxtacribrilina corbicula* | (O'Donoghue & O'Donoghue, 1923) | *Juxtacribrilina* | Cribrilinidae |
| *Kausiaria* sp. | | *Kausiaria* | Otionellidae |
| *Labioporella crenulata* | (Levinsen, 1909) | *Labioporella* | Steginoporellidae |
| *Labioporella spatulata* | Harmer, 1926 | *Labioporella* | Steginoporellidae |
| *Lacerna hosteensis* | Jullien, 1888 | *Lacerna* | Lacernidae |
| *Lacerna watersi* | Hayward & Thorpe, 1989 | *Lacerna* | Lacernidae |
| *Lacrimula burrowsi* | Cook, 1966 | *Lacrimula* | Batoporidae |
| *Lacrimula crassa* | Di Martino, Taylor, Cotton & Pearson, 2017 | *Lacrimula* | Batoporidae |
| *Lacrimula kilwaensis* | Di Martino, Taylor, Cotton & Pearson, 2017 | *Lacrimula* | Batoporidae |
| *Lageneschara lyrulata* | (Calvet, 1909) | *Lageneschara* | Romancheinidae |
| *Lagenicella* n. sp. | | *Lagenicella* | Teuchoporidae |
| *Laminopora jellyae* | (Levinsen, 1909) | *Laminopora* | Adeonidae |
| *Lanceopora* sp. | | *Lanceopora* | Lanceoporidae |
| *Leiosellina edax* | Canu & Bassler, 1923 | *Leiosellina* | Celleporidae |
| *Lunulites bouei* | Lea, 1833 | *Lunulites* | Lunulitidae |
| *Lunulites pyripora* | Canu, 1922 | *Lunulites* | Lunulitidae |
| *Macropora levinseni* | Brown, 1952 | *Macropora* | Macroporidae |
| *Macropora nodulosa* | Gordon & Taylor, 2008 | *Macropora* | Macroporidae |
| *Macropora centralis* | MacGillivray, 1895 | *Macropora* | Macroporidae |
| *Mamillopora* sp. | | *Mamillopora* | Mamilloporidae |
| *Manzonella* sp. | | *Manzonella* | Microporidae |
| *Melicerita (Melicerita) charlesworthii* | Morris, 1843 | *Melicerita* | Cellariidae |
| *Melicerita* cf. *flabellifera* | Hayward & Winson, 1994 | *Melicerita* | Cellariidae |
| *Melicerita flabellifera* | Hayward & Winson, 1994 | *Melicerita* | Cellariidae |
| *Melicerita latilaminata* | Melicerita latilaminata | *Melicerita* | Cellariidae |
| *Membranipora* cf. *ovalis* | (d'Orbigny, 1852) | *Membranipora* | Membraniporidae |
| *Membranipora membranacea* | (Linnaeus, 1767) | *Membranipora* | Membraniporidae |
| *Membranipora* spp. | | *Membranipora* | Membraniporidae |
| *"Membraniporella" robusta* | Lagaaij, 1952 | *Membraniporella* | Cribrilinidae |
| *Membraniporopsis* sp. | | *Membraniporopsis* | Sinoflustridae |
| *Menipea vectifera* | Harmer, 1923 | *Menipea* | Candidae |
| *Metrarabdotos chipolanum* | (Cheetham, 1968) | *Metrarabdotos* | Metrarabdotosidae |
| *Metrarabdotos moniliferum* | (Milne Edwards, 1836) | *Metrarabdotos* | Metrarabdotosidae |
| *Metroperiella reversa* | (Ulrich & Bassler, 1904) | *Metroperiella* | Bitectiporidae |
| *Metroperiella* sp. 1 | | *Metroperiella* | Bitectiporidae |
| *Micropora brevissima* | Waters, 1904 | *Micropora* | Microporidae |
| *Micropora rimulata* | Canu & Bassler, 1929 | *Micropora* | Microporidae |
| *Micropora* spp. | | *Micropora* | Microporidae |
| *Micropora stellata* | Di Martino, Taylor & Portell, 2019 | *Micropora* | Microporidae |
| *Microporella* aff. *californica* | Busk, 1856 | *Microporella* | Microporellidae |
| *Microporella* aff. *ciliata* | (Pallas, 1766) | *Microporella* | Microporellidae |
| *Microporella* aff. *elegans* | Suwa & Mawatari, 1998 | *Microporella* | Microporellidae |
| *Microporella* aff. *speculum* | Brown, 1952 | *Microporella* | Microporellidae |
| *Microporella* aff. *stellata* | (Verrill, 1879) | *Microporella* | Microporellidae |
| *Microporella agonistes* | Gordon, 1984 | *Microporella* | Microporellidae |
| *Microporella arctica* | Norman, 1903 | *Microporella* | Microporellidae |
| *Microporella californica* | Busk, 1856 | *Microporella* | Microporellidae |
| *Microporella* cf. *formosa* | Suwa & Mawatari, 1998 | *Microporella* | Microporellidae |
| *Microporella* cf. *genisii* | (Audouin, 1826) | *Microporella* | Microporellidae |
| *Microporella* cf. *lunifera* | (Haswell, 1881) | *Microporella* | Microporellidae |
| *Microporella* cf. *orientalis* | Harmer, 1957 | *Microporella* | Microporellidae |
| *Microporella* cf. *pontifica* | Osburn, 1952 | *Microporella* | Microporellidae |
| *Microporella ciliata* | (Pallas, 1766) | *Microporella* | Microporellidae |

| | | | |
|---|---|---|---|
| *Microporella coronata* | (Audouin, 1826) | *Microporella* | Microporellidae |
| *Microporella cribrosa* | Osburn, 1952 | *Microporella* | Microporellidae |
| *Microporella discors* | Uttley & Bullivant, 1972 | *Microporella* | Microporellidae |
| *Microporella gibbosula* | Canu & Bassler, 1930 | *Microporella* | Microporellidae |
| *Microporella harmeri* | Hayward, 1988 | *Microporella* | Microporellidae |
| *Microporella hyadesi* | (Jullien, 1888) | *Microporella* | Microporellidae |
| *Microporella intermedia* | Livingstone, 1929 | *Microporella* | Microporellidae |
| *Microporella lingulata* | Di Martino, Taylor & Gordon, 2020 | *Microporella* | Microporellidae |
| *Microporella maldiviensis* | Harmelin, Ostrovsky, Cáceres-Chamizo & Sanner, 2011 | *Microporella* | Microporellidae |
| *Microporella modesta* | Di Martino, Taylor & Gordon, 2020 | *Microporella* | Microporellidae |
| *Microporella morrisiana* | (Busk, 1859) | *Microporella* | Microporellidae |
| *Microporella ordo* | Brown, 1952 | *Microporella* | Microporellidae |
| *Microporella ordoides* | Di Martino, Taylor & Gordon, 2020 | *Microporella* | Microporellidae |
| *Microporella orientalis* | Harmer, 1957 | *Microporella* | Microporellidae |
| *Microporella rusti* | Di Martino, Taylor, Gordon & Liow, 2017 | *Microporella* | Microporellidae |
| *Microporella sarasotaensis* | Di Martino, Taylor & Portell, 2019 | *Microporella* | Microporellidae |
| *Microporella* spp. | | *Microporella* | Microporellidae |
| *Microporella speculum* | Brown, 1952 | *Microporella* | Microporellidae |
| *Microporella stellata* | (Verrill, 1879) | *Microporella* | Microporellidae |
| *Microporella stenoporta* | Hayward & Taylor, 1984 | *Microporella* | Microporellidae |
| *Microporella svalbardensis* | Kuklinski & Hayward, 2004 | *Microporella* | Microporellidae |
| *Microporella tamiamiensis* | Di Martino, Taylor & Portell, 2019 | *Microporella* | Microporellidae |
| *Microporella tanyae* | Di Martino, Taylor & Gordon, 2020 | *Microporella* | Microporellidae |
| *Microporella trigonellata* | Suwa & Mawatari, 1998 | *Microporella* | Microporellidae |
| *Microporella umbonata* | Hincks, 1883 | *Microporella* | Microporellidae |
| *Microporella ventricosa* | Canu & Bassler, 1929 | *Microporella* | Microporellidae |
| *Microporella vibraculifera* | Hincks, 1883 | *Microporella* | Microporellidae |
| *Mobunula bicuspis* | (Hincks, 1883) | *Mobunula* | Petraliellidae |
| *Mollia amoena* | Gordon, 1986 | *Mollia* | Microporidae |
| *Mollia* cf. *mauritiana* | (Kirkpatrick, 1888) | *Mollia* | Microporidae |
| *Mollia multijuncta* | (Waters, 1879) | *Mollia* | Microporidae |
| *Monoporella* n. sp. | | *Monoporella* | Monoporellidae |
| *Monoporella* sp. 1 | | *Monoporella* | Monoporellidae |
| *Mucropetraliella* n. sp. | | *Mucropetraliella* | Petraliellidae |
| *Mucropetraliella* sp. 1 | | *Mucropetraliella* | Petraliellidae |
| *Mucropetraliella thenardii* | (Audouin, 1826) | *Mucropetraliella* | Petraliellidae |
| *Myriapora bugei* | d'Hondt, 1975 | *Myriapora* | Myriaporidae |
| *Myriapora truncata* | (Pallas, 1766) | *Myriapora* | Myriaporidae |
| *Odontoporella* sp. | | *Odontoporella* | Hippoporidridae |
| *Omanipora pilleri* | Berning & Ostrovsky, 2011 | *Omanipora* | Celleporidae |
| *Onychocella* cf. *angulosa* | (Reuss, 1847) | *Onychocella* | Onychocellidae |
| *Otionella perforata* | Canu & Bassler, 1917 | *Otionella* | Otionellidae |
| *Otionellina squamosa* | (Tenison Woods, 1880) | *Otionellina* | Otionellidae |
| *Parafigularia taylori* | (Kukliński & Barnes, 2009) | *Parafigularia* | Cribrilinidae |
| *Parantropora* cf. *laguncula* | (Canu & Bassler, 1929) | *Parantropora* | Antroporidae |
| *Parantropora laguncula* | (Canu & Bassler, 1929) | *Parantropora* | Antroporidae |
| *Parasmittina alanbanneri* | Soule & Soule, 1973 | *Parasmittina* | Smittinidae |
| *Parasmittina aotea* | (Brown, 1952) | *Parasmittina* | Smittinidae |
| *Parasmittina* cf. *raigii* | (Audouin, 1826) | *Parasmittina* | Smittinidae |
| *Parasmittina* cf. *serrula* | Soule & Soule, 1973 | *Parasmittina* | Smittinidae |
| *Parasmittina* cf. *spondylicola* | Harmelin, Bitar & Zibrowius, 2009 | *Parasmittina* | Smittinidae |
| *Parasmittina egyptiaca* | (Waters, 1909) | *Parasmittina* | Smittinidae |
| *Parasmittina latiavicularia* | (Kirkpatrick, 1888) | *Parasmittina* | Smittinidae |
| *Parasmittina* n. sp. | | *Parasmittina* | Smittinidae |
| *Parasmittina parsevalii* | (Audouin, 1826) | *Parasmittina* | Smittinidae |
| *Parasmittina protecta* | (Thornely, 1905) | *Parasmittina* | Smittinidae |
| *Parasmittina raigii* | (Audouin, 1826) | *Parasmittina* | Smittinidae |
| *Parasmittina soulesi* | Scholz & Cusi, 1993 | *Parasmittina* | Smittinidae |
| *Parasmittina* spp. | | *Parasmittina* | Smittinidae |
| *Parasmittina spondylicola* | Harmelin, Bitar & Zibrowius, 2009 | *Parasmittina* | Smittinidae |
| *Parasmittina tropica* | (Waters, 1909) | *Parasmittina* | Smittinidae |
| *Parellisina mirellae* | Di Martino & Taylor, 2014 | *Parellisina* | Calloporidae |
| *Parkermavella* sp. | | *Parkermavella* | Parkermavella |
| *Patsyella acanthodes* | Gordon, 1982 | *Patsyella* | Chaperiidae |
| *Pentapora fascialis* | (Pallas, 1766) | *Pentapora* | Bitectiporidae |
| *Petalostegus* cf. *harmeri* | Gordon & d'Hondt 1991 | *Petalostegus* | Petalostegidae |
| *Petalostegus harmerii* | Gordon & d'Hondt 1991 | *Petalostegus* | Petalostegidae |
| *Petasosella* sp. | | *Petasosella* | Otionellidae |
| *Petralia undata* | MacGillivray, 1869 | *Petralia* | Petraliidae |
| *Petraliella* sp. | | *Petraliella* | Petraliellidae |
| *Phidolopora* sp. | | *Phidolopora* | Phidoloporidae |
| *Phonicosia* n. sp. | | *Phonicosia* | Lacernidae |
| *Phonicosia circinata* | (MacGillivray 1869) | *Phonicosia* | Lacernidae |

| | | | |
|---|---|---|---|
| *Plesiocleidochasma laterale* | (Harmer, 1957) | *Plesiocleidochasma* | Phidoloporidae |
| *Plesiocleidochasma* cf. *vestitum* | (Canu & Bassler, 1923) | *Plesiocleidochasma* | Phidoloporidae |
| *Plesiocleidochasma* sp. 1 | | *Plesiocleidochasma* | Inapplicable |
| *Pleurocodonellina signata* | (Waters, 1889) | *Pleurocodonellina* | Smittinidae |
| *Pleurocodonellina* sp. 1 | | *Pleurocodonellina* | Smittinidae |
| *Pleurocodonellina* n. sp. | | *Pleurocodonellina* | Phidoloporidae |
| *Pleuromucrum gorgonense* | (Hastings, 1930) | *Pleuromucrum* | Phidoloporidae |
| *Pleuromucrum saucatsense* | Vigneaux, 1949 | *Pleuromucrum* | Phidoloporidae |
| *Pleuromucrum* sp. 1 | | *Pleuromucrum* | Phidoloporidae |
| *Porella belli* | (Dawson, 1859) | *Porella* | Bryocryptellidae |
| *Porella* cf. *hanaishiensis* | Hayami, 1975 | *Porella* | Bryocryptellidae |
| *Porella columbiana* | O'Donoghue & O'Donoghue, 1923 | *Porella* | Bryocryptellidae |
| *Porella hanaishiensis* | Hayami, 1975 | *Porella* | Bryocryptellidae |
| *Poricella mucronata* | (Smitt, 1873) | *Poricella* | Arachnopusiidae |
| *Poricella robusta* | (Hincks, 1884) | *Poricella* | Arachnopusiidae |
| *Poricella* sp. 1 | | *Poricella* | Arachnopusiidae |
| *Poricella spathulata* | (Canu & Bassler, 1929) | *Poricella* | Arachnopusiidae |
| *Powellitheca waipukurensis* | (Waters, 1887) | *Powellitheca* | Powellithecidae |
| *Prenantia firmata* | (Waters, 1887) | *Prenantia* | Smittinidae |
| *Puellina* aff. *voigti* | Ristedt, 1985 | *Puellina* | Cribrilinidae |
| *Puellina bifida* | (d'Hondt, 1970) | *Puellina* | Cribrilinidae |
| *Puellina caesia* | Dick, Grischenko & Mawatari, 2005 | *Puellina* | Cribrilinidae |
| *Puellina* cf. *harmeri* | Ristedt, 1985 | *Puellina* | Cribrilinidae |
| *Puellina* cf. *innominata* | (Couch, 1844) | *Puellina* | Cribrilinidae |
| *Puellina harmeri* | Ristedt, 1985 | *Puellina* | Cribrilinidae |
| *Puellina innominata* | (Couch, 1844) | *Puellina* | Cribrilinidae |
| *Puellina quadrispinosa* | Di Martino, Taylor & Portell, 2017 | *Puellina* | Cribrilinidae |
| *Puellina scripta* | (Reuss, 1848) | *Puellina* | Cribrilinidae |
| *Puellina* spp. | | *Puellina* | Cribrilinidae |
| *Pyriporella* sp. | | *Pyriporella* | Calloporidae |
| *Ralepria conforma* | Hayward, 1991 | *Ralepria* | Lacernidae |
| *Reptadeonella brasiliensis* | Almeida, Souza, Sanner & Vieira, 2015 | *Reptadeonella* | Adeonidae |
| *Reptadeonella heckeli* | (Reuss, 1848) | *Reptadeonella* | Adeonidae |
| *Reptadeonella* spp. | | *Reptadeonella* | Adeonidae |
| *Reptadeonella violacea* | (Johnston, 1847) | *Reptadeonella* | Adeonidae |
| *Retelepralia macmonagleae* | Di Martino & Taylor, 2012 | *Retelepralia* | Cheiloporinidae |
| *Reteporella fenestrata* | (Powell, 1967) | *Reteporella* | Phidoloporidae |
| *Reteporella* sp. 1 | | *Reteporella* | Phidoloporidae |
| *Rhynchozoon ferocula* | Hayward, 1988 | *Rhynchozoon* | Phidoloporidae |
| *Rhynchozoon larreyi* | (Audouin, 1826) | *Rhynchozoon* | Phidoloporidae |
| *Rhynchozoon* sp. 1 | | *Rhynchozoon* | Phidoloporidae |
| *Robertsonidra argentea* | (Hincks, 1881) | *Robertsonidra* | Robertsonidridae |
| *Robertsonidra* cf. *argentea* | (Hincks, 1881) | *Robertsonidra* | Robertsonidridae |
| *Robertsonidra praecipua* | Hayward & Ryland, 1995 | *Robertsonidra* | Robertsonidridae |
| *Schizolepraliella nancyae* | Di Martino, Taylor & Portell, 2017 | *Schizolepraliella* | Incertae sedis |
| *Schizomavella (Calvetomavella) phterocopa* | Reverter-Gil, Berning & Souto, 2015 | *Schizomavella (Calvetomavella)* | Bitectiporidae |
| *Schizomavella (Schizomavella) auriculata* | (Hassall, 1842) | *Schizomavella* | Bitectiporidae |
| *Schizomavella (Schizomavella) richardi* | (Calvet, 1903) | *Schizomavella* | Bitectiporidae |
| *Schizomavella* cf. *incompta* | Hayward, 1988 | *Schizomavella* | Bitectiporidae |
| *Schizomavella incompta* | Hayward, 1988 | *Schizomavella* | Bitectiporidae |
| *Schizomavella* spp. | | *Schizomavella* | Bitectiporidae |
| *Schizoporella subsinuosa* | Canu, 1914 | *Schizoporella* | Schizoporellidae |
| *Schizoporella* sp. 1 | | Inapplicable | Inapplicable |
| *Schizoporella unicornis* | (Johnston in Wood, 1844) | *Schizoporella* | Schizoporellidae |
| *Schizosmittina melanobater* | Gordon, 1989 | *Schizosmittina* | Bitectiporidae |
| *Schizosmittina planovicellata* | Vigneaux, 1949 | *Schizosmittina* | Bitectiporidae |
| *Schizotheca* sp. | | *Schizotheca* | Phidoloporidae |
| *Scorpiodinipora* cf. *costulata* | (Canu & Bassler, 1929) | *Scorpiodinipora* | Hippoporidridae |
| *Selenaria maculata* | (Busk, 1852) | *Selenaria* | Selenariidae |
| *Selenaria occidenta* | Cook & Chimonides, 1985 | *Selenaria* | Selenariidae |
| *Selenaria semilunaria* | Bock & Cook, 1999 | *Selenaria* | Selenariidae |
| *Selenaria vera* | Bock & Cook, 1999 | *Selenaria* | Selenariidae |
| *Septentriopora karasi* | Kukliński & Taylor, 2006 | *Septentriopora* | Calloporidae |
| *Setosellina* cf. *constricta* | Harmer, 1926 | *Setosellina* | Heliodomidae |
| *Setosellina constricta* | Harmer, 1926 | *Setosellina* | Heliodomidae |
| *Setosinella perfluxa* | Di Martino & Taylor, 2012 | *Setosinella* | Pyrisinellidae |
| *Setosinella prolifica* | Canu & Bassler, 1933 | *Setosinella* | Pyrisinellidae |
| *Smittina anecdota* | Hayward & Thorpe, 1990 | *Smittina* | Smittinidae |
| *Smittina diffidentia* | Hayward & Thorpe, 1989 | *Smittina* | Smittinidae |
| *Smittina directa* | (Waters, 1904) | *Smittina* | Smittinidae |
| *Smittina glebula* | Hayward & Thorpe, 1990 | *Smittina* | Smittinidae |
| *Smittina hanaishiensis* | Hayami, 1975 | *Smittina* | Smittinidae |
| *Smittina rogickae* | Hayward & Taylor, 1984 | *Smittina* | Smittinidae |

| | | | |
|---|---|---|---|
| *Smittina* sp. 1 | | *Smittina* | Smittinidae |
| *Smittipora* aff. *cordiformis* | Harmer, 1926 | *Smittipora* | Onychocellidae |
| *Smittipora* cf. *cordiformis* | Harmer, 1926 | *Smittipora* | Onychocellidae |
| *Smittipora cordiformis* | Harmer, 1926 | *Smittipora* | Onychocellidae |
| *Smittipora* sp. | | *Smittipora* | Onychocellidae |
| *Smittoidea malleata* | Hayward & Thorpe, 1989 | *Smittoidea* | Smittinidae |
| *Smittoidea maunganuiensis* | (Waters, 1906) | *Smittoidea* | Smittinidae |
| *Smittoidea pugiuncula* | Hayward & Thorpe, 1989 | *Smittoidea* | Smittinidae |
| *Spiniflabellum jacksoni* | Di Martino, Taylor & Portell, 2017 | *Spiniflabellum* | Cribrilinidae |
| *Spiniflabellum laurae* | Di Martino, Taylor & Portell, 2019 | *Spiniflabellum* | Cribrilinidae |
| *Steginoporella cornuta* | Osburn, 1950 | *Steginoporella* | Steginoporellidae |
| *Steginoporella cucullata* | (Reuss, 1848) | *Steginoporella* | Steginoporellidae |
| *Steginoporella discors* | Gordon, Voje & Taylor, 2017 | *Steginoporella* | Steginoporellidae |
| *Steginoporella magnifica* | Harmer, 1900 | *Steginoporella* | Steginoporellidae |
| *Steginoporella neozelanica* | (Busk, 1861) | *Steginoporella* | Steginoporellidae |
| *Steginoporella rhomboidalis* | Canu & Lecointre, 1927 | *Steginoporella* | Steginoporellidae |
| *Steginoporella* spp. | | *Steginoporella* | Steginoporellidae |
| *Steginoporella turbarens* | Canu & Lecointre, 1927 | *Steginoporella* | Steginoporellidae |
| *Stephanollina vorax* | (Canu & Bassler, 1923) | *Stephanollina* | Sfeniellidae |
| *Stylopoma* cf. *duboisii* | (Audouin, 1826) | *Stylopoma* | Schizoporellidae |
| *Stylopoma* cf. *novum* | Tilbrook, 2001 | *Stylopoma* | Schizoporellidae |
| *Stylopoma* cf. *vilaensis* | Tilbrook, 2001 | *Stylopoma* | Schizoporellidae |
| *Stylopoma farleyensis* | Di Martino, Taylor & Portell, 2017 | *Stylopoma* | Schizoporellidae |
| *Stylopoma palmula* | Tilbrook, 2001 | *Stylopoma* | Schizoporellidae |
| *Stylopoma* spp. | | *Stylopoma* | Schizoporellidae |
| *Stylopoma vilaensis* | Tilbrook, 2001 | *Stylopoma* | Schizoporellidae |
| *Stylopoma viride* | (Thornely, 1905) | *Stylopoma* | Schizoporellidae |
| *Taylorus arcuatus* | (Gordon, 2014) | *Taylorius* | Escharinidae |
| *Taylorus spinosus* | (Gordon, 2014) | *Taylorius* | Escharinidae |
| *Taylorus waiparaensis* | (Brown, 1952) | *Taylorius* | Escharinidae |
| *Tegella aquilirostris* | (O'Donoghue & O'Donoghue, 1923) | *Tegella* | Calloporidae |
| *Tegella cassidata* | (O'Donoghue & O'Donoghue, 1923) | *Tegella* | Calloporidae |
| *Thairopora* sp. | | *Thairopora* | Thalamoporellidae |
| *Thalamoporella biperforata* | Canu & Bassler, 1923 | *Thalamoporella* | Thalamoporellidae |
| *Thalamoporella californica* | (Levinsen, 1909) | *Thalamoporella* | Thalamoporellidae |
| *Thalamoporella* cf. *rozieri* | (Audouin, 1826) | *Thalamoporella* | Thalamoporellidae |
| *Thalamoporella rasmuhammadi* | Soule, Soule & Chaney, 1999 | *Thalamoporella* | Thalamoporellidae |
| *Thalamoporella rozieri* | (Audouin, 1826) | *Thalamoporella* | Thalamoporellidae |
| *Thalamoporella sibogae* | Soule, Soule & Chaney, 1992 | *Thalamoporella* | Thalamoporellidae |
| *Thalamoporella* sp. | | *Thalamoporella* | Thalamoporellidae |
| *Thalamoporella spinosa* | Chaney, Soule & Soule, 1989 | *Thalamoporella* | Thalamoporellidae |
| *Thornelya* sp. 1 | | *Thornelya* | Hippopodinidae |
| *Thornelya* sp. 2 | | *Thornelya* | Hippopodinidae |
| *Toretocheilum turbinatu* | Hayward, 1995 | *Toretocheilum* | Escharinidae |
| *Tornipora canui* | (Brydone, 1906) | *Tornipora* | Onychocellidae |
| *Torquatella* sp. | | *Torquatella* | Celleporidae |
| *Trematooecia* cf. *turrita* | (Smitt, 1873) | *Trematooecia* | Colatooeciidae |
| *Triphyllozoon* sp. 1 | | *Triphyllozoon* | Phidoloporidae |
| *Triphyllozoon* sp. 2 | | *Triphyllozoon* | Phidoloporidae |
| *Triphyllozoon* sp. 3 | | *Triphyllozoon* | Phidoloporidae |
| *Triporula stellata* | (Smitt, 1873) | *Triporula* | Exechonellidae |
| *Trypostega* sp. | | *Trypostega* | Trypostegidae |
| *Trypostega venusta* | (Norman, 1864) | *Trypostega* | Trypostegidae |
| *Turbicellepora bernardi* | (Audouin, 1826) | *Turbicellepora* | Celleporidae |
| *Turbicellepora redoutei* | (Audouin, 1826) | *Turbicellepora* | Celleporidae |
| *Turbicellepora smitti* | (Kluge, 1962) | *Turbicellepora* | Celleporidae |
| *Umbonula littoralis* | Hastings, 1944 | *Umbonula* | Umbonulidae |
| *Valdemunitella fraudatrix* | Gordon, 1986 | *Valdemunitella* | Calloporidae |
| *Valdemunitella huttoni* | (Brown, 1952) | *Valdemunitella* | Calloporidae |
| *Valdemunitella pyrula* | (Hincks, 1881) | *Valdemunitella* | Calloporidae |
| *Vibracellina capillaria* | Canu & Bassler, 1917 | *Vibracellina* | Cupuladriidae |
| *Vibracellina viator* | Canu & Bassler, 1929 | *Vibracellina* | Cupuladriidae |
| *Villicharixa strigosa* | (Uttley, 1951) | *Villicharixa* | Electridae |
| *Watersipora* cf. *subovoidea* | (d'Orbigny, 1852) | *Watersipora* | Watersiporidae |
| *Watersipora* cf. *subtorquata* | (d'Orbigny, 1852) | *Watersipora* | Watersiporidae |
| *Watersipora* sp. | | *Watersipora* | Watersiporidae |
| *Watersipora subovoidea* | (d'Orbigny, 1852) | *Watersipora* | Watersiporidae |
| *Xenogma rhomboidale* | (Powell, 1967) | *Xenogma* | Buffonellodidae |
| *Yrbozoon conspicuum* | (Powell, 1967) | *Yrbozoon* | Cleidochasmatidae |
| *Zeuglopora* sp. | | *Zeuglopora* | Conescharellinidae |
